# RNA virus genomes from centuries- to millennia-old Adélie penguin mummies

**DOI:** 10.64898/2025.12.17.693957

**Authors:** Tjorven Hinzke, Chris Lauber, Katharina J. Hoff, Jennifer Klunk, Madeline Tapson, Emilio Mármol-Sánchez, Marc A. Suchard, Philippe Lemey, Steven D. Emslie, Sébastien Calvignac-Spencer

## Abstract

Direct studies of long-term RNA virus evolution are largely limited to chemically-fixed specimens from museums collected over the past two centuries. Detecting genomic traces of RNA viruses in older, buried remains is considered highly unlikely. The cold, dry conditions of Antarctica may represent an exception. Here, natural mummification of penguins and seals—animals that form large colonies where RNA viruses circulate—is common, potentially facilitating RNA virus genome recovery. Here, we show that Adélie penguin (*Pygoscelis adeliae*) mummies, ranging in age from recent to nearly two millennia, indeed contain ancient viral genome fragments. Metatranscriptomic analyses yielded near-complete genome sequences of a picornavirus (*Megrivirus epengu*) and a rotavirus D (*Rotavirus deltagastroenteritidis*) from relatively recent specimens. We further retrieved rotavirus D sequences from a 280-year-old individual and a near-complete rotavirus G (*Rotavirus gammagastroenteritidis*) genome from a 2000-year-old one. These findings pave the way to direct studies of RNA virus evolution across millennia.

## Introduction

Fossils are essential for reconstructing the deep evolutionary history of cellular organisms.^1^ By offering irreplaceable insights into past biological diversity, they provide anchors to infer key evolutionary mechanisms and the pathways that led to contemporary life forms. Viruses do not leave a physical fossil record; however, the ability to sequence ancient viral DNA has already begun to transform the study of DNA virus evolutionn.^2,3^ Extending such approaches to RNA viruses, which can exhibit extraordinarily high rates of genetic change^4–6^ and which have often experienced recent genetic bottlenecks,^7^ holds similar promise. Ancient RNA virus genomes could provide unprecedented insights into the evolutionary space these viruses once explored, including unique opportunities to reveal key genetic factors enabling cross-species transmission. Such knowledge is critical for understanding how RNA viruses interact with other biological entities in diverse ecosystems, but also for preparing for new outbreaks and pandemics.

Some insights into ancient RNA virus evolution have come from endogenous viral elements (EVEs), viral genomes or genome fragments that were integrated into host germline DNA and inherited over potentially long evolutionary timescales.^8,9^ EVEs are broadly regarded as an evolving fossil record, and have accumulated mutations since integration. Therefore, they do not provide unaltered information about past RNA virus genetic diversity. In addition, most studied endogenization events are extremely ancient, often dating back millions of years. At the other end of the temporal spectrum, metatranscriptomic analyses of chemically fixed pathological specimens have enabled direct study of RNA virus genomes from the last two centuries.^10–12^ This leaves a vast chronological gap in which direct recovery of vertebrate RNA virus genomes has generally be considered nearly impossible^13^ (see ^14–17^ for plant and insect RNA viruses).

Here, we asked whether it is possible to retrieve, under the most favorable conditions, centuries- or even millennia-old RNA virus genetic material from well-preserved and mummified vertebrate remains. Host RNA has been successfully recovered from ancient permafrost-preserved samples and even from museum specimens stored at room temperature.^18–22^ If host RNA can survive for such extended periods, could viral RNA, often present in large quantities during infection, also be detectable? We explored this possibility in remains with exceptional potential: mummified Adélie penguins (*Pygoscelis adeliae*) from Antarctica.

In this extremely cold and dry region, ancient nucleic acids can indeed be remarkably well preserved, as shown by the recent recovery of ancient DNA (aDNA) from thousand-year-old sediment with minimal damage.^23^ Mummification occurs naturally in Antarctica, and researchers regularly discover Adélie penguin mummies at sites where colonies have persisted for centuries or even millennia.^24,25^ Some mummies could stem from individuals that died of infections with RNA viruses, as a number of these viruses are known to circulate today in the densely populated colonies.^26–31^ The mummies may provide insights into globally relevant RNA viruses, which have already reached Antarctica, such as the recently arrived highly pathogenic avian influenza virus H5N1.^32^

## Results and Discussion

### RNA virus genomes in modern Adélie penguin mummies

We began by analyzing four modern Adélie penguin chick mummies, collected opportunistically on Jan 14, 2016 at an active breeding site at Cape Hallett, Ross Sea (CH1, CH2, CH3, CH4; Figure 1). We did not attempt radiocarbon dating, as their recent origin was clear from their fresh appearance and their location on the surface of an active penguin colony. The specimens received no specific treatment, were transported in plastic bags and stored upon arrival at −20°C until sampling for analysis in 2023. We sampled small pieces of tissue from the throat, cloaca, and lateral thorax (targeting lung tissue between the ribs), since these anatomical sites offer good chances of RNA virus detection in modern birds. We extracted total nucleic acids from all samples, and prepared metagenomic libraries for CH3 and CH4 and metatranscriptomic libraries for all specimens, which we sequenced on Element Biosciences (RNA from CH1 and CH2) and Illumina (DNA and RNA from CH3 and CH4) platforms. On average, we obtained 14.3 ± 3.6·10^6^ (mean ± standard deviation) metagenomic and 19.2 ± 10.0·10^6^ (Illumina) or 31.4 ± 20.5·10^6^ (Element Biosciences) metatranscriptomic read pairs per sample (Table 1, Supplementary Tables 1, 2), which were used for downstream host and viral screening.

**Figure 1.**
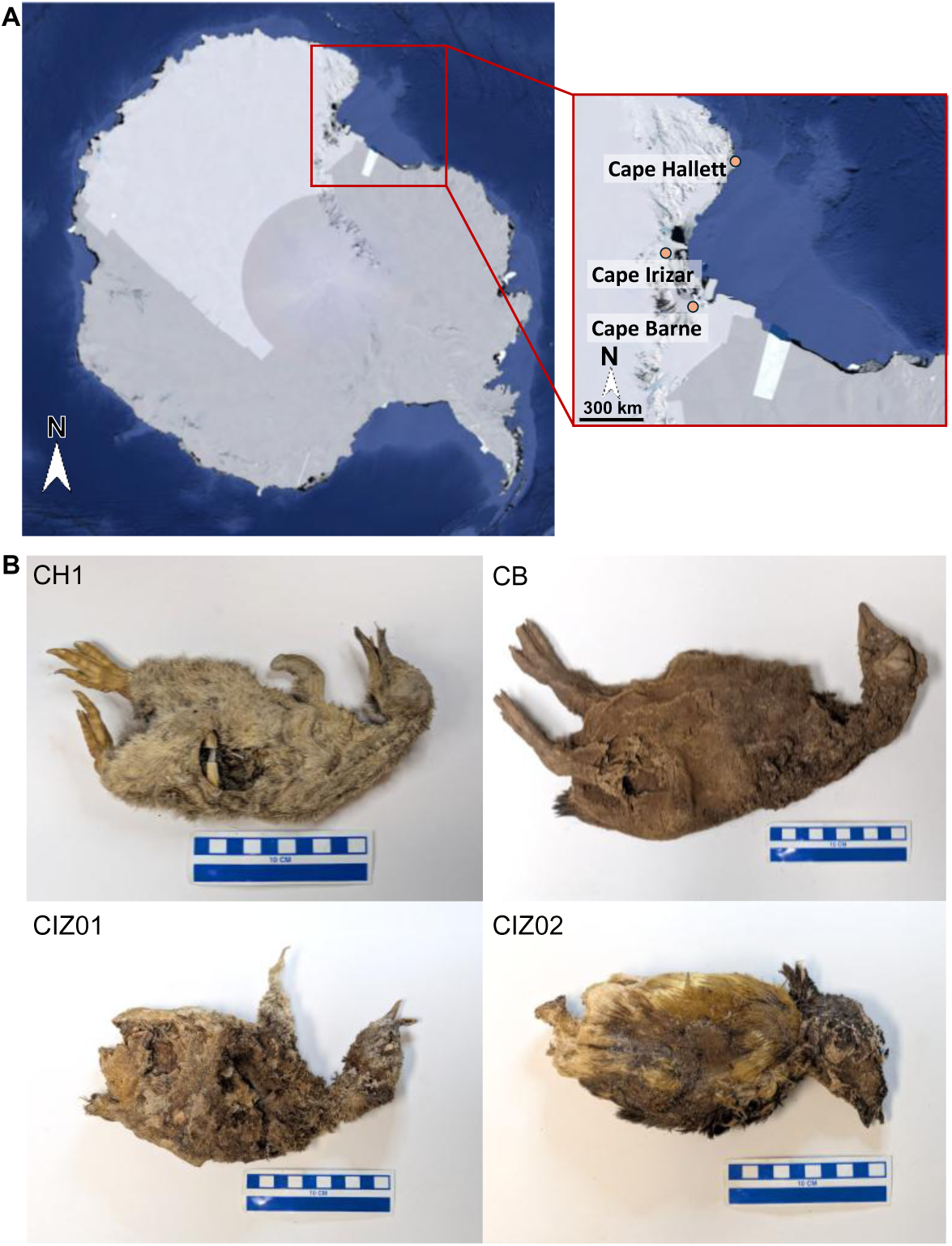
Adélie penguin mummies from Antarctica with viral remains. (A) Map of Antarctica with inset (red rectangle) for the Ross Sea region showing precise locations of Adélie penguin sites where chick mummies were recovered (Google Earth, 14.02.2025). (B) Adélie penguin chick mummies in which RNA virus genomes were found.

**Table 1.** Specimens, their estimated age, sampling locations, storage after sampling, and sequencing results. All specimens were collected in January 2016, except for CB, which was collected in January 2001. See main text for details on estimated age and Supplementary Tables 1 and 2 for sequencing results. RT: room temperature, EB: Element Biosciences, I: Illumina.

| | Estimated age | Sampling location | Storage | Metatranscriptome read pairs mean ± standard deviation $\cdot 10^6$ (sequencing platform) | Metagenome read pairs mean ± standard deviation $\cdot 10^6$ (all Illumina) |
| --- | --- | --- | --- | --- | --- |
| CH1 | after 2000 | Cape Hallett | -20 °C | 19.5±21.6 (EB) | NA |
| CH2 | after 2000 | Cape Hallett | -20 °C | 43.3 ± 9.7 (EB) | NA |
| CH3 | after 2000 | Cape Hallett | -20 °C | 17.0 ± 6.9 (I) | 13.0 ± 4.4 |
| CH4 | after 2000 | Cape Hallett | -20 °C | 21.1 ± 11.6 (I) | 15.7 ± 1.6 |
| CB | 280 cal. yr. BP | Cape Barne | dry, RT | 11.6 ± 4.4 (EB) | NA |
| CIZ01 | 735 cal. yr. BP | Cape Irizar | dry, RT | 32.2 ± 22.9 (I) | 15.6 ± 3.3 |
| CIZ02 | 1913 cal. yr. BP | Cape Irizar | dry, RT | 23.8 ± 16.6 (I) | 13.1 ± 2.4 |

We used metagenomic reads to assemble almost complete Adélie penguin mitochondrial genomes for CH3 and CH4 (97.9 and 98.0% unambiguous nucleotides in the final consensus as compared to the reference mitogenome NC_021137.1) Trimmed and quality-filtered DNA fragments had a mean length of 97 (CH3) and 86 (CH4) bp and showed minimal damage, with terminal misincorporation frequencies mostly below 0.01 (Supplementary Figure 1).

We also mapped paired metatranscriptomic reads to mitogenome consensus sequences created from metatranscriptomic reads, to compare DNA vs. RNA preservation. Mean trimmed and quality-filtered RNA fragment lengths were 112 (CH1), 84 (CH2), 68 (CH3), and 83 (CH4) nt, and terminal deamination frequencies were mostly below 0.01, aside from CH1, with relatively few reads mapping to the incomplete consensus and some frequencies above 0.1 (Supplementary Figure 2). Generally, both DNA and RNA appeared to be well-preserved in most of these specimens.

We screened metatranscriptomic reads for the presence of viruses by profile Hidden Markov Model (pHMM)-based homology search and *de novo* sequence assembly. The resulting contigs were compared to virus and host reference sequence databases obtained from the NCBI to validate putative viral hits. This analysis indicated that two specimens (CH1, CH4) were positive for at least one RNA virus. We detected numerous megrivirus (genus *Megrivirus*, family *Picornaviridae*) reads in thorax, cloacal, and/or throat samples from CH1 and CH4 (6.4–1004.6 reads per million reads (rpm), Supplementary Table 3). Low numbers of megrivirus reads were detected in other samples mapping to final assemblies (22 reads in CH4 cloacal samples, 8 reads in CH2 cloacal samples), which may indicate lower viral RNA abundance at other anatomical sites and/or limited bleed-through during sequencing.

Megriviruses form non-enveloped, icosahedral viral particles containing a single-stranded (ss) positive-sense RNA genome of a length of 9040-9840 nt.^33^ The data enabled near-complete genome reconstruction of megrivirus genomes from both penguin mummies (96.6–98.8 % coverage relative to the closest reference sequences MN453780.1, MN453781.1, and NC_039004.1). The recovered genomes only differed at 0.58% of positions (Supplementary Table 4) and therefore likely originated from the same virus species. The respective reads showed somewhat higher damage as compared to the host sequences, but terminal misincorporation frequencies were still mostly below 0.03 (Supplementary Figure 3). Maximum-likelihood phylogenetic analyses confirmed that these two viruses were closest relatives (Figure 2), and that their nearest known relative is another megrivirus previously detected in Adélie penguins and ornithogenic soil at an Adélie penguin colony (*Megrivirus epengu*, NC_039004,^33,34^ see also Supplementary Results). Megriviruses have frequently been detected in Antarctic penguin species,^31,34–36^ and one of them was found to be abundant in yellow-eyed penguin chicks (*Megadyptes antipodes*) that developed diphteric stomatitis, but to be absent from healthy adults.^37^ The detection of *Megrivirus epengu* in tissues of CH1 and CH4 provides a first indication that recently mummified specimens allow for RNA virus detection and reconstruction of RNA virus genomes.

**Figure 2.**
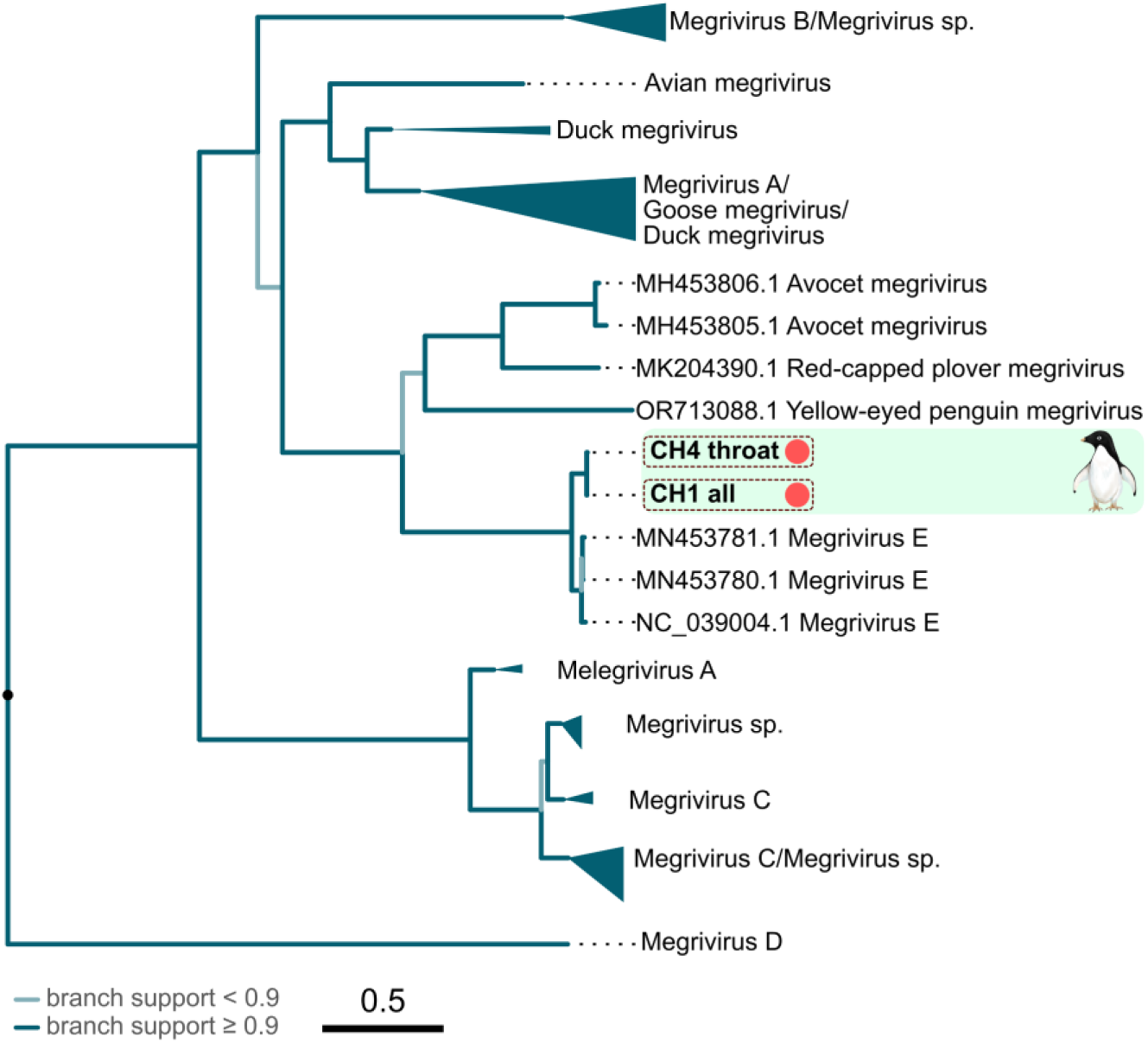
Adélie penguin mummy megriviruses are closest related to *Megrivirus epengu*. Megrivirus sequences obtained in this study are highlighted. Branch colors reflect bootstrap support based on IQ-TREE’s UFBoot algorithm with 2000 bootstrap replicates.

The detection of rotavirus reads in the cloacal samples of CH4 (195.4-203.8 rpm, Supplementary Table 3) further corroborated the potential for viral RNA recovery from Adélie penguin mummies. Rotaviruses (family *Sedoreoviridae*) form non-enveloped, icosahedral viral particles that enclose a genome of 11 linear double-stranded (ds) RNA segments with a total genome length of ∼18,600 bp. They are often associated with gastrointestinal disease in young mammals and birds.^38^ We reconstructed a near-complete rotavirus genome from CH4, recovering sequences covering between 81.5 and 112.8% of the 11 genomic segments (Figure 3A; in some cases sequences obtained by *de novo* assembly were longer than those of the reference genome). Phylogenetic analyses showed that this virus was most closely related to rotavirus D (aside from another virus also detected in this study, see below), a bird-specific viral species (*Rotavirus deltagastoenteritidis*; Figure 3B,C, Supplementary Figure 4-13), in all segment products but segment 1 (pairwise distance (p-distance) 24.4%-56.6%, Supplementary Tables 5-15). The amino acid product of segment 1 was most closely related to Mawson virus (p-distance 2.9%), which was identified from contemporary Adélie penguins and for which two VP1 sequences are available.^31^ The next-closest relative to CH4 in VP1 is again a rotavirus D (p-distance: 32.4%; Figure 3C and Supplementary Table 5). These results indicate that years- to decades-old mummified remains from Antarctica can yield well-preserved RNA genomes from different viral taxa (terminal misincorporation frequency mostly below 0.002; Supplementary Figure 14). The detection of three RNA viruses in four individuals, including one co-infection (CH4), also suggests that a sizeable fraction of now-mummified Adélie penguin chicks were infected with RNA viruses at the time of their death.

**Figure 3.**
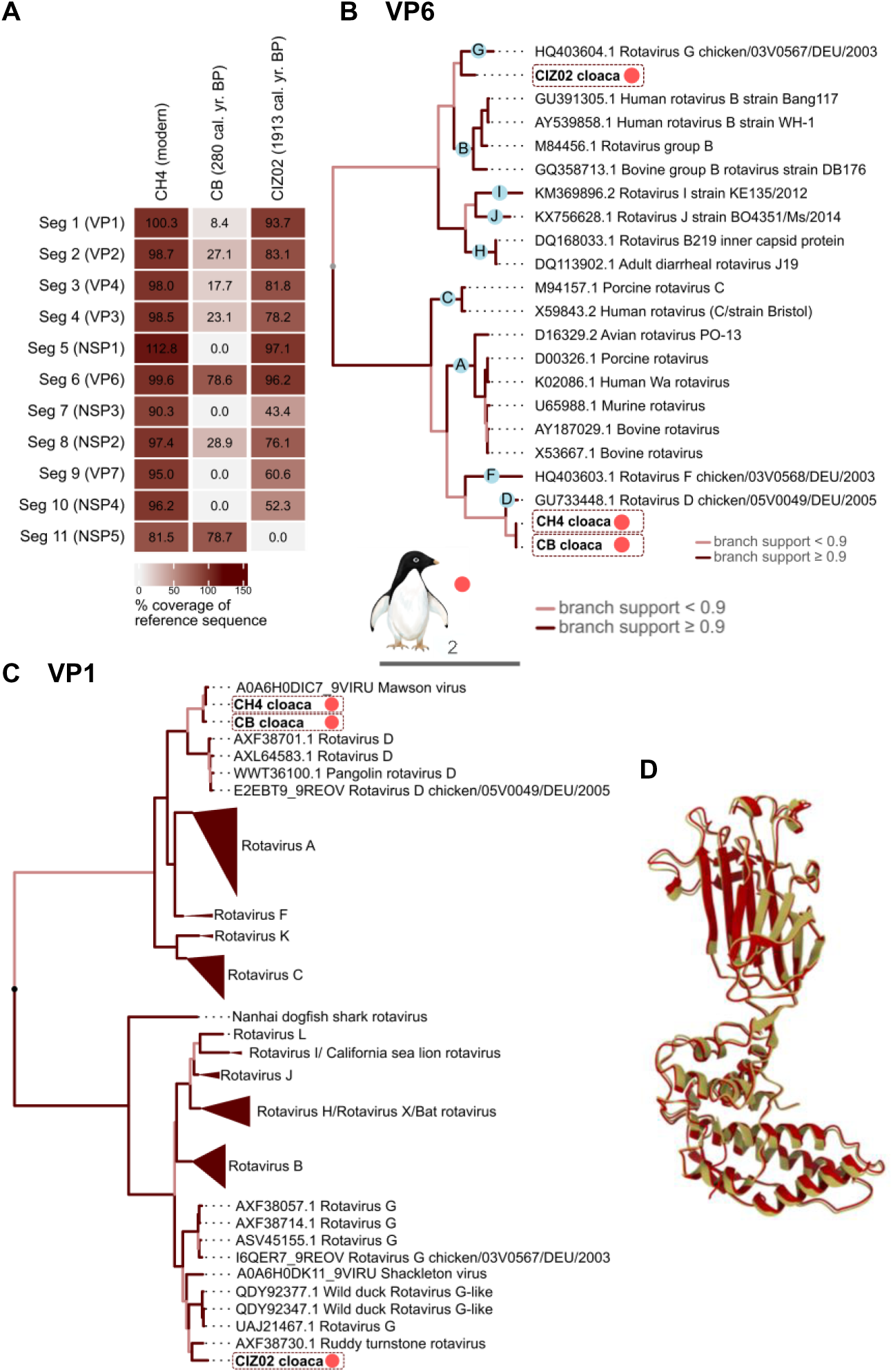
Rotavirus sequences of Adélie penguin mummies are closely related to rotavirus D and G. (A) Percentage coverage of rotavirus nucleotide sequences obtained in this study in relation to their closest ICTV relatives (CIZ02: rotavirus G chicken/03V0567/DEU/2003, CH4, CB: rotavirus D chicken/05V0049/DEU/2005). A percentage coverage >100% indicates that sequences obtained by de novo assembly in this study are longer than those of the respective reference serotype. We did not identify NSP6 as gene product of segment 11. Seg: segment. (B) Phylogeny of rotavirus VP6 obtained in this study (red dots) and VP6 of ICTV reference strains. (C) Phylogeny of rotavirus VP1 obtained in this study (red dots) and published VP1. Phylogenies based on amino acid sequences. Note that the legend pertains to Panels B and C. Branch support based on IQ-TREE’s UFBoot algorithm with 2000 bootstrap replicates. (D) Reconstructed 2000-year-old rotavirus G VP6 structure. Alphafold models of CIZ02 rotavirus VP6 (red), superimposed on the nearest relative ruddy turnstone rotavirus (AXF38736.1, beige).

### RNA virus genomes in Adélie penguin mummies up to 2000 years old

Following our encouraging results obtained from modern Adélie penguin mummies, we analyzed three much more ancient mummies collected at two abandoned Adélie penguin colonies at Cape Barne and Cape Irizar^24,39^ (Figure 1). Cape Barne was last occupied by breeding Adélie penguins at about 300-400 calibrated years before present (cal. yr. BP; Emslie, unpubl. data), while Cape Irizar was last occupied at about 685-1100 cal. yr. BP.^39^ In good agreement with these estimates of colony occupancy, the mummies were dated with radiocarbon analysis to a mean 280 cal. yr. BP (CB), 735 cal. yr. BP (CIZ01) and 1913 cal. yr. BP (CIZ02; see Supplementary Table 16 for values of replicate measurements). The specimens had been collected in 2001 (CB) and 2016 (CIZ01, CIZ02) and were kept at ambient temperature in a metal storage cabinet until 2023. They were sampled and processed in the same way as previously described, and we sequenced metagenomics libraries for CIZ01 and CIZ02 and metatranscriptomic libraries for all specimens. We generated on average 14.4 ± 3.3·10^6^ metagenomic and 27.3 ± 19.9·10^6^ (Illumina) or 11.6 ± 4.4·10^6^ (Element Biosciences) metatranscriptomic read pairs per sample (Table 1, Supplementary Tables 1,2).

Using metagenomic reads, we reconstructed a nearly complete mitogenome from CIZ02, and a highly fragmented one from CIZ01 (CIZ02: 97.8%, CIZ01: 21.8%). The CIZ02 sequence differed from those of the more recent specimens with p-distances of 1.8% (CIZ01), 1.9% (CH4), and 5.9% (CH3; Supplementary Table 17). Three of the four mitogenomes here (CH4, CIZ01, CIZ02) apparently belong to the same group and one (CH3) to another (Supplementary Figure 15). We initially aimed to corroborate the radiocarbon dating using temporal signal in Adélie penguin mitochondrial sequences. However, the temporal signal was almost exclusively supported by two 37,600 to 44,000 years-old sequences, and we did not find a significant correlation of genetic divergence and age when only younger specimens were included in our analyses, preventing any meaningful attempt at sequence-based age estimation^40,41^ (Supplementary Figure 16–18). DNA fragments mapping to the mitogenomes had a mean length of 102 bp (CIZ01) and 89 bp (CIZ02) and exhibited low damage (none for CIZ01; terminal misincorporation frequency <0.01 for CIZ02; Supplementary Figure 19). Additionally, we mapped metatranscriptomic reads to the reference mitogenome, observing 0.2-96.4 metatranscriptomic rpm in these ancient specimens, compared to 0.6-1481.4 rpm in the more recent ones (for Illumina; for Element Biosciences: 27.3-496.3 vs. 0.02-880.1 rpm; Supplementary Table S2), with slightly higher misincorporation frequencies as compared to the DNA reads (Supplementary Figure 20). These results show the presence of endogenous nucleic acids in the older mummies.

We screened metatranscriptomic reads of CB, CIZ01, and CIZ02 as described above for RNA viral sequences and detected rotaviruses in the cloacal samples of CB and CIZ02. A second library generated from an independent sample of CIZ02 confirmed the rotavirus detection but only produced very few viral reads (Supplementary Table S3). Viral abundance in CB and CIZ02 was lower than in the cloacal sample from the more recent CH4 (lower than 21 rpm vs around 200 rpm; Supplementary Table S3). This lower value may reflect different viral loads or increased degradation in less well-preserved specimens. While misincorporation frequencies sometimes exceeded 0.05 in rotavirus reads of CB, they were mostly below 0.006 in CIZ02 (Supplementary Figure 21). We were able to reconstruct 7 partial segments for CB (8.4-78.6%), and 10 for CIZ02 (43.4-97.1 %) with mean coverage read depth of 6.0-12.7 and 6.8-30.4-fold per segment. Phylogenetic analyses of all 7 segment products available for the rotavirus from CB showed it to be the closest relative of the virus identified in CH4 (p-distances 0.3-13.9%; only for VP3 the p-distance was artifactually 0% for a rotavirus K due to overlap in only one amino acid, Figure 3B,C; Supplementary Figure 4-6, 8, 10, 13; Supplementary Tables 5-8, 10, 12, 15). In contrast, the rotavirus from CIZ02 was most closely related to the ruddy turnstone rotavirus or other rotavirus G representatives (species *Rotavirus gammagastroenteritidis*) in all segments (Figure 3, Supplementary Figure 4-12; Supplementary Tables 5-14). The phylogeny was largely reflected in the p-distances: for all but products of segments 3 and 5, the closest matches were ruddy turnstone rotavirus or rotavirus G (22.7-51.8%), whereas segments 3 and 5 were most similar to rotavirus B and H (33.7 and 66.9%), respectively. Ruddy turnstones (*Arenaria interpres*) are cosmopolitan shorebirds, but the individual that led to the identification of the rotavirus was sampled in Australia.^42^ Taken together, these analyses identify RNA virus genomes in ancient Adélie penguin mummies that belong to lineages plausibly associated with these hosts.

We subsequently performed more detailed phylogenetic analyses focusing on the higher quality and oldest CIZ02 rotavirus genome. To this end, we assembled a dataset comprising the publicly available sequences most closely related to CIZ02 in VP6 (n=18), one of the more conserved rotavirus segments^43^ for which CIZ02 showed good coverage. As in previous analyses, the closest match was the ruddy turnstone rotavirus. In a maximum likelihood phylogeny, however, CIZ02 did not fall on a noticeably shorter branch, contrary to the expectation that ancient viruses often exhibit measurably shorter branches.^44^ Instead, its branch was even markedly longer than that of the ruddy turnstone virus (Supplementary Figure 22). This pattern is not necessarily surprising given that CIZ02 and the ruddy turnstone virus differed at 22% of amino acid positions in VP6 (Supplementary Table 10), indicating considerable divergence and a deep evolutionary history in a sparsely sampled part of the rotavirus phylogeny. Under such conditions, extensive substitution on long branches can obfuscate the relationship between divergence and sampling time.

To characterize the substitution process and place this divergence on an explicit timescale, we estimated the VP6 codon substitution rate in a heterochronous rotavirus A (*Rotavirus alphagastroenteriditis*) VP6 data set with strong temporal signal.^45^ We confirmed the generalizability of this rate estimate by obtaining highly similar values for rotavirus C (*Rotavirus tritogastroenteridis*).^46^ We then used the rotavirus A estimate as evolutionary rate calibration and modeled potential variation in purifying selection through time in the CIZ02 VP6 data set,^10^ incorporating the radiocarbon age as sampling time for CIZ02. The model supported a declining (log) dN/dS ratio into the past, suggesting stronger long-term purifying selection (posterior mean slope = −0.24 per unit natural logarithm-time, 95% HPD = [−0.38, −0.11]). Structural modeling of VP6 revealed highly similar predicted structures for CIZ02 and the ruddy turnstone rotavirus, consistent with such strong purifying selection acting on the capsid protein (Figure 3D). As expected, the time-heterogeneous selection model also estimated a root age considerably older than the estimate under a time-homogenous model (mean age: 32680 years, 95% HPD = 9637-62854, vs. 12493 years, 95% HPD = 7240-18884).

Finally, to assess whether CIZ02 should be expected to show a shorter root-to-tip distance than the ruddy turnstone virus under this estimated evolutionary process, we simulated segment 6 sequence evolution along the inferred tree and analyzed the root-to-tip distances in 1000 replicates. In 66.3% of simulations, CIZ02 did not have the lowest root-to-tip distance, and in 13% of simulations the CIZ02 root-to-tip distance exceeded that of the ruddy turnstone virus. Thus, the observed branch length of CIZ02 falls within the range expected under a plausible evolutionary model and is not in conflict with its radiocarbon age. More broadly, these results indicate that in deeply divergent and sparsely sampled viral lineages, the absence of detectable branch shortening cannot be taken as evidence against authenticity, as substitution saturation and undersampling can obscure the relationship between divergence and sampling time.

### Frozen archives of RNA virus evolution

Overall, we recovered genomic material from a picornavirus and several rotaviruses in a remarkably small sample of four modern and three ancient Adélie penguin specimens. Authentication is a major challenge in ancient pathogen research, and in this respect it is noteworthy that the viral sequences presented here did not fulfill two frequently applied criteria: they did not display clear age-dependent nucleic acid damage profiles,^47^ and the most ancient virus did not exhibit a shorter branch in phylogenetic analyses. However, low levels of damage in several specimens are consistent with the excellent preservation conditions, as illustrated by comparably low damage on the host side. Limited apparent damage might also reflect the characteristics of our molecular approach. Indeed, a recent comparison of our method with an alternative in the context of RNA virus sequencing from century-old pathology specimens showed that it does not capture existing damage patterns.^48^ In addition, our in-depth exploration of the CIZ02 divergence revealed that, despite its age, a shorter branch is not necessarily expected under realistic substitution processes across deep evolutionary histories and in the absence of more closely related modern relatives. Together with the use of an ancient DNA facility, the known avian host range of the detected viruses (including Adélie penguins), and their recovery from anatomically relevant sites of penguin mummies from long-abandoned colonies, these observations support the authenticity of our findings. We therefore conclude that under these specific circumstances, it is possible to recover RNA virus sequences hundreds to thousands of years old.

Following from this, an open question is the identification of alternative substrates for ancient viral RNA discovery. Frozen marine bird and mammal mummies are unique to Antarctica, but other frozen sources of ancient viral RNA could exist, including glacier ice and permafrost.^49^ In fact, one of the oldest vertebrate RNA virus genomes sequenced to date was recovered from a victim of the 1918 influenza pandemic buried in Alaskan permafrost,^12,50^ and a likely insect-associated partial RNA virus was detected in 700-year-old frozen caribou feces from Siberia.^15^ In the Ross Sea region, Adélie penguin colony sediments up to 6,000 years old have revealed excellent preservation of aDNA from penguins, seals and their prey,^23^ which perhaps also highlights conditions favorable to ancient RNA survival. Assessing such samples for ancient viral RNA could greatly expand the range of suitable substrates.

We believe that our understanding of RNA virus evolution could be profoundly reshaped and refined by hundreds- to thousands-years-old viral genomes from polar environments. That these exceptional natural archives are under immediate threat from climate change underscores the urgency of collecting potentially suitable samples as well as identifying existing collections amenable to viral RNA detection.

## Limitations of the study

While our results are encouraging, it remains unclear whether they predict the broad survival of RNA virus genomes in ancient mummified remains from Antarctica. We detected only rotaviruses in the ancient specimens. Rotaviruses have double-stranded RNA genomes, which are likely to have higher stability than single-stranded RNA genomes. Moreover, they form triple-layered icosahedral particles^51^ that are particularly resistant to environmental degradation.^52,53^ The long-term preservation of rotavirus nucleic acids, or related viruses with double-stranded RNA genomes and viral particles of comparable stability, may therefore be an exception among RNA viruses. This question can only be answered by analyzing much larger sample sets. Nevertheless, our detection of a megrivirus – a single-stranded RNA virus – in a recent mummy where a rotavirus was also present suggests that single-stranded RNA viruses may also leave long-lasting traces in similar specimens.

## Methods

### Sampling locations

Adélie penguin chick mummies were collected from abandoned penguin colonies at two locations in the Ross Sea region, Antarctica: Cape Barne (CB), Ross Island (−77.58S, 166.24E), and Cape Irizar (CIZ), Lamplugh Island (–75.55S, 162.95E). The penguin mummy from CB was found approximately 10 cm below the surface during excavations of an abandoned colony on 26 January 2001. The chick mummies at CIZ (CIZ01 and CIZ02) were both found on the surface of abandoned penguin colonies on 28 January 2016. All three mummies were wrapped in aluminum foil and placed in clean plastic bags, then transported to the laboratory and stored in metal geology cabinets at room temperature. Cape Hallett (CH; −72.31S, 170.21E) is a large active Adélie penguin colony that was visited on 14 January 2016. The surface of the colony had dozens of fresh and older, mummified carcasses of penguin chicks scattered throughout the area. Four of the older carcasses were collected and returned to the laboratory where they were kept frozen at −20° C until analysis.

### Tissue sampling

A total of seven Adélie penguin mummies, three of which were ancient, were sampled for this study (Table 1). Samples were taken from the regions of the throat/trachea, the lateral lung/rib (thorax) and the cloacae by using sterile scissors and a scalpel cleaned with 10% bleach solution to remove 1-3 g of tissue. The tissues were wrapped in aluminum foil and stored in sterile plastic Whirl-Pak bags until analysis.

### Radiocarbon dating of penguin mummies

Skin samples from mummies or preserved modern species was used. All radiocarbon dates were completed at either the University of Georgia Center for Applied Isotope Studies AMS laboratory (UGAMS) or the Woods Hole National Ocean Sciences AMS Facility (NOSAMS) and are registered with their lab numbers.

UGAMS: Acetone was used to remove fat from the penguin skin samples, and samples were subsequently rinsed in several steps at room temperature with HCl, deionized water, 0.1 N NaOH, again with HCl, deionized water, and dried. Samples were combusted at 900 °C in evacuated ampoules with CuO. CO_2_ was cryogenically purified and converted to graphite.^54^ 14C/13C ratios were measured in a CAIS 0.5 MeV accelerator mass spectrometer (AMS). Sample ratios were compared to that from Oxalic Acid I (SRM 4990).

NOSAMS: Penguin skin samples were placed in 5-10 ml of 1.2 M HCl, followed by a water bath for 1 h after which they were centrifuged at 2300 rpm for 3 min. Samples were rinsed with Milli-Q water, then placed in 0.5 M NaOH and a water bath for 1 h after which they were centrifuged again before decanting the supernatant. This step was repeated up to 10 times until the solution was clear, then the samples were rinsed with Milli-Q water and 5-10 ml of 1.2 M HCl was added, followed by a water bath for 1 h. The samples were then centrifuged, decanted, and rinsed with Milli-Q water until neutralized to a pH of 5-6. Each sample was filtered onto a quartz fiber filter that had been pre-baked at 650° C for 1 h and rinsed with Milli-Q water, then dried in a drying oven at 50 °C for 24-36 h. 14C/13C ratios were measured at the NOSAMS accelerator mass spectrometer (AMS) with sample ratios compared to that from NBS Oxalic Acid I (NIST-SRM-4990).

All radiocarbon dates were corrected and calibrated for the marine carbon reservoir effect using Calib^55^ 8.2 and the Marine20 calibration curve^56^ with a ΔR value of 609 +/- 137 years^57^ to provide a 2-sigma calibration range in cal. yr. BP (Supplementary Table 16).

### Sequencing

Adélie penguin mummy samples were prepared in two different batches for sequencing (Supplementary Table 18). All samples were handled in an ancient DNA clean room. Samples were cut into smaller pieces using disposable sterile tools or non-disposable tools sterilized with a 6% bleach solution. All surfaces were cleaned with 6% bleach before, between, and after handling samples. Four consecutive 24-hour rounds of demineralization and digestion were performed at room temperature (25°C) with shaking at 1000 rpm in 500 µL of buffer, following Schwarz et al. (2009).^58^ Samples containing bone (a subset of the thorax samples) received alternating rounds of demineralization and digestion (4 total rounds), samples containing no bone received digestion only (4 total rounds). Extraction was performed for all samples following the protocol outlined by Dabney et al. (2013)^59^ with the substitution of High Pure Viral Nucleic Acid Large Volume column (Roche) for the spin column step. Total DNA mass was quantified via Qubit 1X dsDNA High Sensitivity (HS; ThermoFisher Scientific).

For batch 1, half of the extracted volume was used for DNA library preparation. The extract was split in two again in order to prepare two independent DNA libraries per DNA extract. Up to 5 ng of extracted DNA was taken into a single-stranded DNA library preparation method (SRSLY PicoPlus, Claret Bioscience) following the manufacturer’s protocol. Libraries were amplified for 14 cycles and quantified via Qubit 1X dsDNA HS. The other half of the extracted volume was DNase treated with Turbo-DNA free (ThermoFisher Scientific) and purified with SPRI beads at a 3X ratio to generate an RNA extract. Total RNA mass was quantified via Qubit HS RNA assay (ThermoFisher Scientific). The RNA extract was split in two to prepare two independent libraries per extract. RNA library preparation was performed using the KAPA RNA Hyper Prep kit (Roche) following the manufacturer’s instructions for degraded samples. For indexing amplification, 14 cycles of amplification were performed. Samples were quantified via the Qubit 1X dsDNA HS. Based on initial quantifications and the calculated mass required for sequencing, samples were reamplified for 8-32 cycles and quantified again. Libraries were visualized via TapeStation 4200 (Agilent) on a D1000 High Sensitivity screen tape at 1 ng µL^-1^. Samples that showed evidence of bubbling (bimodal distribution) received three cycles of amplification to recondition the libraries. Samples that showed the presence of adapter artifacts (a sharp peak at ∼134 bp) were subjected to artifact removal. Samples requiring artifact removal were pooled at equimolar ratios into four pools with samples of similar morphology and size selected to remove adapter dimer via Blue Pippin (Sage Science). Libraries and size-selected pooled libraries were combined into the final sequencing pool at approximately equimolar ratios. The final sequencing pool was visualized via TapeStation as above. The samples were sequenced on an Illumina NovaSeq 6000 platform using v1.5 chemistry on a portion of an S4 PE150 flowcell targeting a yield of approximately 6 Gbp per library. Data were demultiplexed and subsequently used for analysis as described below. The size-selected pool containing one or both RNA libraries from the samples CIZ01 throat, CIZ01 thorax, CIZ02 throat (two libraries), CIZ02 cloaca (two libraries), and CH4 throat was sequenced again on the same platform to approximately 6 Gbp per library.

Batch 2 was processed using the same methods as for the first batch, with the following modifications: No DNA libraries were prepared, so 80% of the total extracted volume (up to 3 µg) was used for DNase treatment. RNA was visualized via TapeStation 4200 (Agilent) after DNase treatment and bead-based purification with a High Sensitivity RNA tape. Libraries were index amplified for 12 cycles. Samples CIZ01 cloaca and CIZ02 cloaca (four libraries from two extracts) were sequenced as above. The remainder of the samples were sequenced on the AVITI platform (Element Biosciences) on two PE150 Cloudbreak high-throughput flowcells.

## Data analysis

### Virus identification and viral genome sequence assembly

To identify viral sequences in the sequencing data and to generate consensus contigs of viral genome sequences, two workflows were integrated:

i. This two-stage workflow consists of a sensitive sequence homology-based virus identification step followed by target viral sequence assembly. Raw reads were adapter- and quality-trimmed using fastp^60^ version 0.23.2, converted to Fasta format using Seqtk version 1.0-r82-dirty (https://github.com/lh3/seqtk), and *in silico* translated in all six reading frames using EMBOSS transeq^61^ version 6.5.7.0. For initial identification of RNA viral sequences, we screened the translated polypeptide sequences for sequence homology to the hallmark protein of RNA viruses, the RNA-dependent RNA polymerase (RdRp), using a sensitive, profile Hidden Markov Model (pHMM)-based approach through hmmsearch from the HMMER package^62^ version 3.4. We included pHMMs of RdRps from a comprehensive set of RNA viruses, as described previously.^63,64^ We considered hits with hmmsearch E-values of 10 or smaller, which were further validated by sequence assembly in the second workflow stage. In this second workflow stage, we sought to reconstruct as much sequence information as possible from the potentially fragmented or incompletely sequenced genomes or genome segments for the viral hits obtained in the first stage. To do so, we first compared the trimmed sequencing reads to reference amino acid sequences of a virus group of interest (e.g., reoviruses or picornaviruses) using Diamond blastx^65^ version 2.0.13.151 with an E-value cutoff of 1e-5. The resulting hits were subsequently assembled into contigs using CAP3^66^ with parameters –o 20 –h 99. We then sought to iteratively extend these contigs. We used them as query in a MMseqs2 easy-search^67^ version 5f8735872e189991a743f7ed03e7c9d1f7a78855 analysis to identify additional reads that align to the contig termini. The current set of contigs and the newly identified reads were again assembled using CAP3, potentially resulting in elongated contigs. This process was repeated 10 times. Two final rounds of CAP3 were conducted to merge any remaining overlapping contigs/singlets.
ii. Raw reads were adapter- and quality-trimmed and deduplicated using fastp^60^ 0.23.4. PolyG and polyX sequences, as well as low-complexity sequences (threshold 15) and sequence duplicates (accuracy 6) were removed. Remaining reads were assembled *de novo* into contigs using megahit^68^ v1.2.9. These contigs were searched against viral RefSeq nucleotides (BLAST^69^ 2.15.0+) with discontinuous megablast (sequences downloaded from NCBI 11.01.2024, default E-value 10), and against viral RefSeq proteins (Diamond^65^ 2.1.8 blast in very-senstitive mode, default E-value 0.001; sequences downloaded from NCBI 01.03.2024). Initial hits against viral reference sequences were re-blasted against the NCBI nucleotide (nt) database (downloaded 08.01.2024) and NCBI non-redundant (nr) protein database (downloaded 28.02.2024) to remove false-positive hits, and to only retain viral sequences. The remaining viral contigs of interest were concatenated with read-based identification data resulting from workflow ii (see below). Viral hits were filtered from blast results using a custom R^70^ script (R version 4.3.1), with the packages seqinr^71^ 4.2-36 and dplyr^72^ 1.1.4. For read mapping against contigs, we used bbmap.sh from the BBTools suite^73^ (version 39.06). Additionally, we used samtools^74^ 1.19.2 for sam to bam conversion and sorting.

Subsequently, the contigs and reads of interest from the two workflows were concatenated, de-duplicated and containments integrated on a 100% identity level using clumpify.sh from the BBTools suite. For subsequent remapping, the trimmed and filtered reads from workflow part (i) were used. All reads mapping to the contigs of interest were retained. Resulting reads were assembled with SPAdes^75^ v3.15.5, using --sc -k 21,33,55,77,99,127. Reads were remapped onto these contigs and filtered for mapq>25. The pipeline was implemented in snakemake^76^ 8.3.2.

The final consensus of rotavirus sequences was generated in Geneious v. 2023.0.4 (https://www.geneious.com; highest quality (50%), call N if coverage < 3, call Sanger Heterozygotes > 50 %). For the Picornaviridae, as the *de novo* assembly clearly showed the presence of megrivirus E, we subsequently assembled virus sequences from all CH1 samples and CH4 throat by mapping quality- and adapter-trimmed reads against the megrivirus E NCBI reference genome (NC_039004.1) with bbmap.sh from BBTools, calling the consensi with iVar (https://github.com/andersen-lab/ivar) version 1.4.2, and subsequently using these consensi for an additional round of mapping and consensus generation. After calculating orthoANI scores (https://www.ezbiocloud.net/tools/ani) for the CH1 megrivirus consensus sequences, which were all > 99%, we re-assembled one megrivirus consensus sequence for all three CH1 samples.

### Annotation of viral consensus sequences

Viral consensus sequences were searched against the nr and nt databases (see above), as well as custom *Picornaviridae* (01.07.2024) and *Rotavirus* (01.07.2024) protein (NCBI and UniProt) and nucleotide (NCBI) sequence databases (clustered at 100% with CD-Hit^77^ version 4.8.1). Sequences which generated a majority of top 3 hits for *Picornaviridae* or *Rotavirus* were retained. Annotations for Megrivirus/ other *Picornaviridae* and *Rotavirus* segments were likewise generated by majority vote of the top 3 hits.

### Viral phylogeny

For phylogenetic analysis of rotavirus sequences, we used an amino acid sequence-based approach, given the high divergence of rotaviral sequences. We translated nucleotide contig sequences with all six possible reading frames, using EMBOSS^61^ v.6.6.0.0 transeq. Resulting potential amino acid sequences were then searched using a discontinuous megablast^69^ and Diamond blast^65^ against the respective rotavirus databases (see above), to find the most likely reading frame. We corrected the reading frame for segment 7 of CH4 within the sequence by adding a gap. For subsequent analyses on amino acid level, sequences before and after stop codons were removed, to leave the most likely open reading frames. For initial phylogenetic analyses, we used rotavirus protein sequences from NCBI (downloaded 19.07.2024) and from Uniprot (downloaded 17.11.2024). These rotavirus sequences were subsetted by excluding sequences with *Homo sapiens* as host or with no host given, and with including ICTV sequences (by name of the virus), RefSeq sequences, and sequences for which an animal name was included in the virus denomination. The resulting sequences were partitioned into viral proteins (for the NCBI data, partitioning was done not only on the often missing segment information, but also by subsetting on the respective text strings in the protein names). Obtained sequences were clustered at 95% using CD-Hit. Alignment of own sequences with the thus obtained references was done using mafft^78^ with mafft-linsi. For CIZ02 VP4 we initially obtained three and for CH4 NSP4 two contigs. After alignment assessment, we retained the most contiguous contig for both and re-calculated the respective alignments. We subsequently manually curated the alignments using Aliview^79^ v. 1.28. Alignments were trimmed with BMGE^80^ v2.0. Phylogenetic trees were reconstructed using IQ-TREE 2^81^ v. 2.0.7 with UFBoot2^82^ (2000 bootstrap replicates) and the ModelFinder^83^ module. For further scaffolding, we aligned our sequences to other rotavirus nucleotide sequences (downloaded from NCBI 19.11.2024, subsetted for rotavirus D or G) of the same group, as identified by our original analysis, and merged contigs where possible (i.e., in the case of segment 6 of CIZ02 rotavirus). All subsequent analysis were performed with these final manually curated sequences. For VP6, we additionally generated phylogenetic trees as above using the rotavirus reference sequences used by the International Committee on the Taxonomy of Viruses (ICTV). We used AlphaFold3^84^ to generate structural models of VP6 from CIZ02 and from ruddy turnstone rotavirus (AXF38736.1). The resulting models were superimposed with ChimeraX^85^ 1.9.

Maximum likelihood (ML) phylogenetic reconstruction for the VP6-encoding segment 6 CIZ02 data set was performed using IQ-TREE 2^81^ under the best-fitting substitution model (the general time-reversible model with discrete gamma-distributed rate variation among sites). To estimate a timed evolutionary history for this data set, we fitted a time-dependent selection model^10^ using BEAST^86^ X. This model uses an epoch-parameterization^87^ to allow the nonsynonymous over synonymous substitution (dN/dS) rate ratio in a codon substitution model to vary through time, effectively accommodating potential long-term purifying selection.^88^ In order to calibrate the codon substitution rate in this model, we estimated this rate based on a heterochronous segment 6 data set for rotavirus A with strong temporal signal.^45^ This yielded a codon substitution rate of 1.79E-03 (9.99E-04,2.75E-03) per site per year. A rotavirus C data set produced highly similar rates of 1.69E-03 (9.42E-04,2.40E-3). We assessed the fit of this model relative to a homogeneous codon substitution model using the posterior estimate for the dN/dS decline rate.^10^ We also attempted to accommodate further rate variation among lineages using branch-specific random effects,^10^ but posterior estimates for the standard deviation of these random effects indicated overparameterization.

To evaluate expectations for root-to-tip divergence as a function of tip age in ML phylogenetic reconstructions of the deep segment 6 evolutionary history, we simulated sequence evolution with πBUSS^89^ according to the time-varying selection process estimated using the above model and reconstructed ML trees. Based on 1000 replicates, we determined the frequency by which CIZ02 had the shortest root-to-tip divergence, and the frequency by which it had shorter root-to-tip divergence compared to its sister lineage.

For Megrivirus alignments, we used as reference all megrivirus sequences downloaded from NCBI Virus (11.10.24), and subsequently filtered for a length >3000 bases and clustered on 99% sequence identity with CD-Hit. We used EMBOSS^61^ v.6.6.0.0 getorf for finding the open reading frame (ORF) per sequence. Sequence alignments were done in Seaview^90^ 5, using Clustal-Omega0^91^ 1.1 on the amino acid level. Alignments were manually curated and exported as nucleic acid. We subsequently filtered alignments using BMGE. We tested for recombination events with RDP5 Beta^92^ 5.75, which did not result in any evidence for recombination events impacting the direct phylogenetic neighborhood of our megrivirus sequences. We then calculated phylogenetic trees with IQ-TREE 2^81^ as for the rotavirus sequences.

Pairwise hamming distances (p-distances) of trimmed alignments were calculated in R^93^ v. 4.5.2 using the package phangorn^94^ 2.12.1 with pairwise gap exclusion.

### Coverage heatmap and phylogenetic tree visualization

Coverage heatmap and tree visualizations were generated in R v. 4.5.2 using the packages ape^95^ 5.8-1, circlize^96^ 0.4.16, ComplexHeatmap^97^ 2.24.1, ggtree^98^ 3.16.3, phytools^99^ 2.4-4, seqinr^71^ 4.2-36, and svglite^100^ 2.2.1. Phylogenetic trees were rerooted for visualization by minimizing variance of root to tip distances using FastRoot^101^ v. 1.5.

### Host RNA analysis

Metatranscriptomes were quality-filtered and trimmed using fastp^60^, and mapped against penguin and human reference genomes (penguin: ASM69910v1, human: GRCh38.p14) using hisat2^102^ v. 2.2.1. Mitochondrial consensus sequences were generated as for DNA (see below).

### Host DNA analysis

Metagenomes of four specimens (CIZ01, CIZ02, CH3, CH4) were quality-filtered and trimmed using fastp^60^ and mapped against penguin and human reference genomes (penguin: ASM69910v1, human: GRCh38.p14) using hisat2 v. 2.2.1.

For generating mitochondrial consensus sequences, the same quality-filtered and trimmed metagenome reads were mapped against the Adélie penguin mitochondrial reference genome (NC_021137.1) using bbmap.sh. Consensi were called in Geneious v. 2023.0.4 using 10x coverage and 95% identity thresholds. Reference Adélie mitogenome sequences for alignment were downloaded from NCBI (23.09.2024) and clustered on 99% identity with CD-Hit. Additionally, ancient Adélie penguin mitochondrial sequences^40,41^ were used for phylogenetic analysis. Pairwise distances were calculated in R 4.5.2 using the package phangorn 2.12.1 with pairwise gap exclusion.

For generating phylogenetic trees and estimating evolutionary distances, we created the following data-subsets:

i. Adélie penguin mitogenome sequences obtained in this study and publicly available mitogenome sequences as outlined above and
ii. a re-analysis of the studies of Lambert et al. (2002)^40^ and Subramanian et al. (2009)^41^ without own sequences.

Mitogenome maximum likelihood phylogenetic trees were generated as were viral trees above.. Treetime 0.11.4 (https://github.com/neherlab/treetime) was used for root-to-tip vs. time regression-based dataset (i) initial tree outlier identification, which were subsequently removed for generating the final tree. For inspecting potential time signals in the mitogenomes’ molecular phylogenies, we used TempEst^103^ v. 1.5.3

### Analysis of sequence damage profiles

We used DamageProfiler^104^ v. 1.0 to assess damage patterns of reads mapped against the respective consensus sequences.

## Supporting information

Supplementary Tables and Consensus sequences

## Resource availability

The sequencing data are available at the European Nucleotide Archive (ENA) at EMBL-EBI, accession number PRJEB97393.

## Acknowledgements

This research was funded by the Deutsche Forschungsgemeinschaft (DFG, German Research Foundation) – Project-ID 531801029 – TRR 410 “WETSCAPES2.0”, and Project-ID 522593772. Field work and sample collection in Antarctica was funded by National Science Foundation Grants OPP 9909274 and ANT 1443386. PL and MAS are partially supported through US National Institutes of Health grant R01 AI153044. PL acknowledges support by the Research Foundation - Flanders (‘Fonds voor Wetenschappelijk Onderzoek - Vlaanderen’, G0D5117N, G0B9317N and G051322N). CL is supported by the Deutsche Forschungsgemeinschaft (DFG, German Research Foundation) under Germany’s Excellence Strategy - EXC 2155 - project number 390874280 and by KA1-Co-02 “CoViPa”, a grant from the Helmholtz Association’s Initiative and Network Fund. We thank the myReads team at Daicel Arbor Biosciences for assistance in sample handling. We thank Dr. Marie-Katherin Zühlke for thoughtful input on the protein structure reconstruction, and Dr. Erin Harvey for valuable feedback on the rotavirus phylogeny.

## Author contributions

T.H. performed analyses, wrote the initial manuscript, and prepared Figures. J.K. and M. T. performed sample preparation and sequencing. C.L. performed sequence analyses. K.J.H. contributed to sequence analyses. E. M.-S. contributed to sequence analyses. P.L. performed phylogenetic analyses. M.A.S. contributed to phylogenetic analyses. S.D.E. collected samples in Antarctica and performed laboratory tissue sampling. S.C.-S. designed and supervised the study, and contributed to sequence analyses and initial manuscript writing. All authors contributed to, reviewed, and edited the manuscript.

## Supplemental Information

Supplementary Text including Supplementary Figures 1-22, Supplementary Tables 17, 18 Supplementary Tables 1-15, 17

Supplementary Data: Consensus sequences of host and viruses

