## Supplementary Text for "RNA virus genomes from centuries- to millennia-old Adélie penguin mummies"

### Supplementary Results

#### Virus species demarcations

We classified the viral sequences detected here according to the International Committee on the Taxonomy of Viruses (ICTV) criteria, while not attempting to give a final delineation of the viral species identified. The nearest relative to the megriviruses detected in Adélie penguin mummies in this study is recognized as the species *Megrivirus epengu* by the ICTV, and the megriviruses detected in CH1 and CH4 samples fulfil the criteria to be considered members of this species, with an overall amino acid divergence of 12.2% (CH1) and 10.7% (CH4) to the megrivirus E1 reference sequence NC_039004.1. The rotaviruses detected in this work likely belong to *Rotavirus gammagastroenteritidis* (rotavirus G; CIZ02) and *Rotavirus deltagastroenteritidis* (rotavirus D; CH4, CB), based on the ICTV cutoff of 53% identity of the VP6 amino acid sequence.[1]

#### Mapping statistics and damage patterns

Ratio of RNA reads mapping to the *P. adeliae* vs. human reference genome was about 0.7-~170x; mostly 1-5x (Supplementary Table 2), and the respective ratio for DNA reads varied between 1.3 > 300x (Supplementary Table 1). There was no clear relation between age of samples and read mapping ratio. Likewise, damage profiles were not clearly correlated with age, i.e., older mummies did not show consistently higher DNA damage (Supplementary Figure 19).

### Supplementary Figures

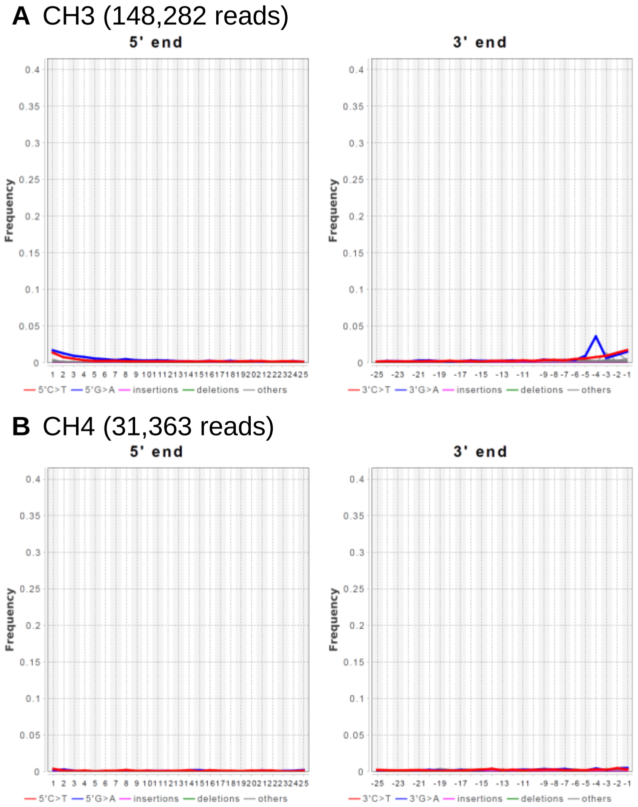


**Supplementary Figure 1. Damage patterns of modern *Pygoscelis adeliae* mummy DNA library reads aligned to their respective consensus sequences**
(A) Adélie penguin mummy CH3.
(B) Adélie penguin mummy CH4.

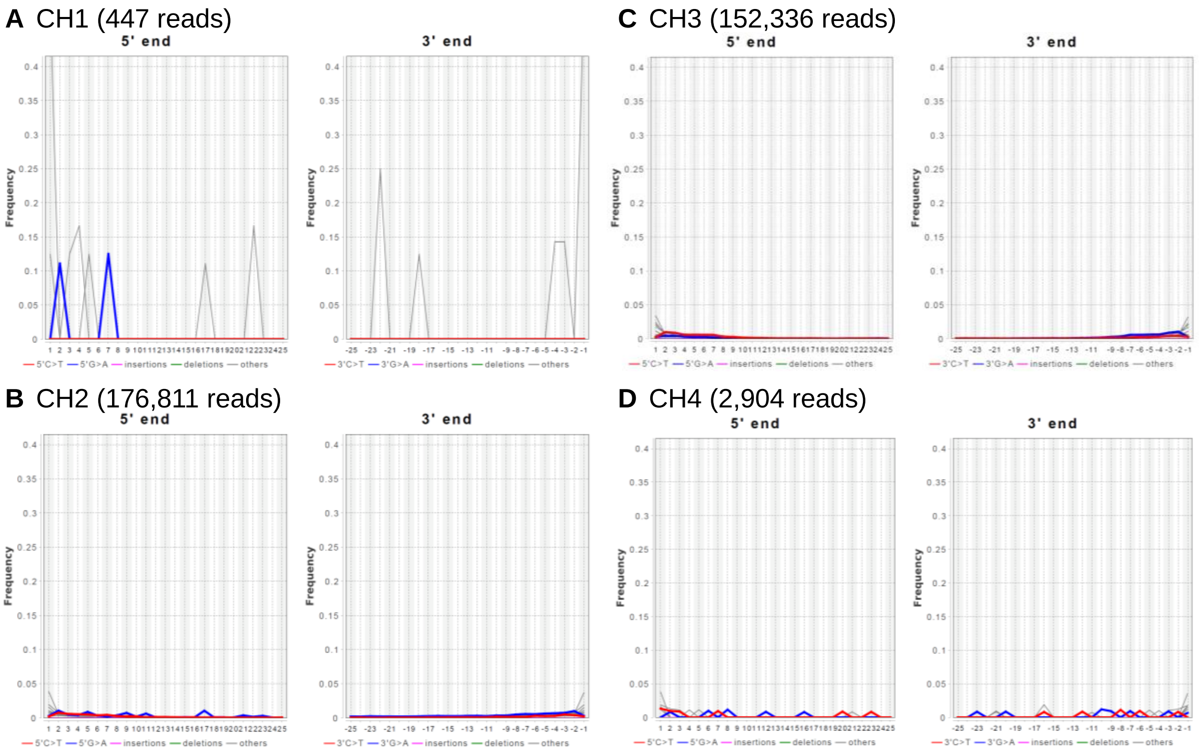


**Supplementary Figure 2. Damage patterns of modern *Pygoscelis adeliae* mummy RNA library reads aligned to their respective RNA consensus sequences**
(A) Adélie penguin mummy CH1.
(B) Adélie penguin mummy CH2.
(C) Adélie penguin mummy CH3.
(D) Adélie penguin mummy CH4.

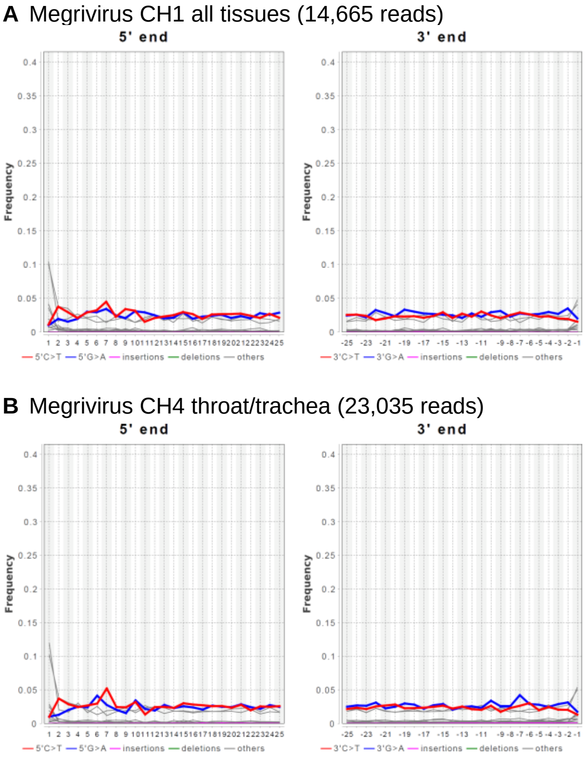


**Supplementary Figure 3. Damage patterns of reads aligned to megrivirus consensus sequences in modern *Pygoscelis adeliae* mummy RNA library reads**
(A) Adélie penguin mummy CH1 (all tissues).
(B) Adélie penguin mummy CH4 (throat/tracheal region).

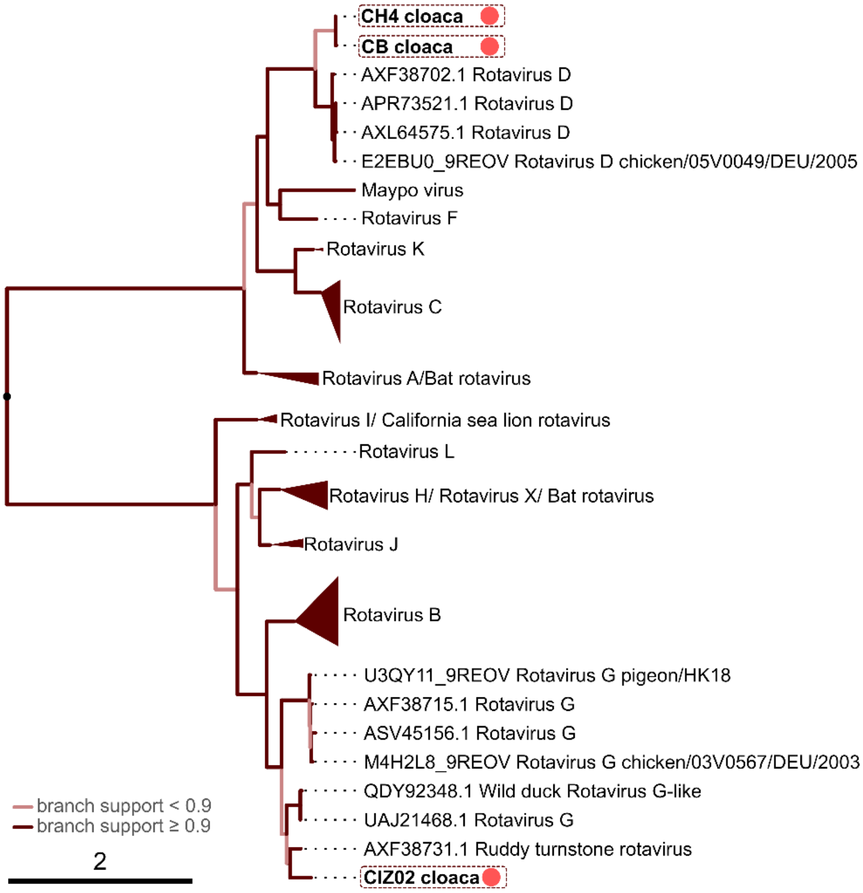


**Supplementary Figure 4. Phylogenetic tree of rotavirus VP2 (product of segment 2) sequences obtained in this study (red dots) and published rotavirus VP2 sequences**
Phylogenies based on amino acid sequences.

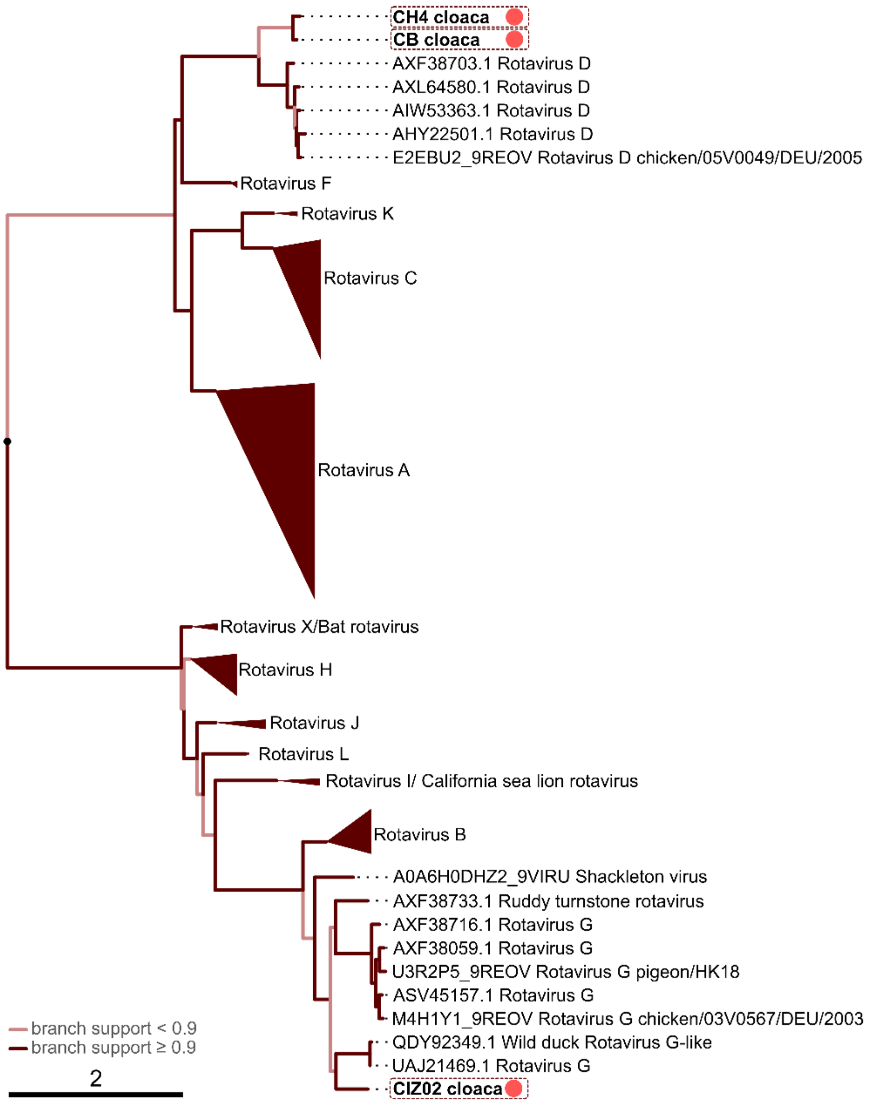


**Supplementary Figure 5. Phylogenetic tree of rotavirus VP3 (product of segment 3) sequences obtained in this study (red dots) and published rotavirus VP3 sequences**

Phylogenies based on amino acid sequences.

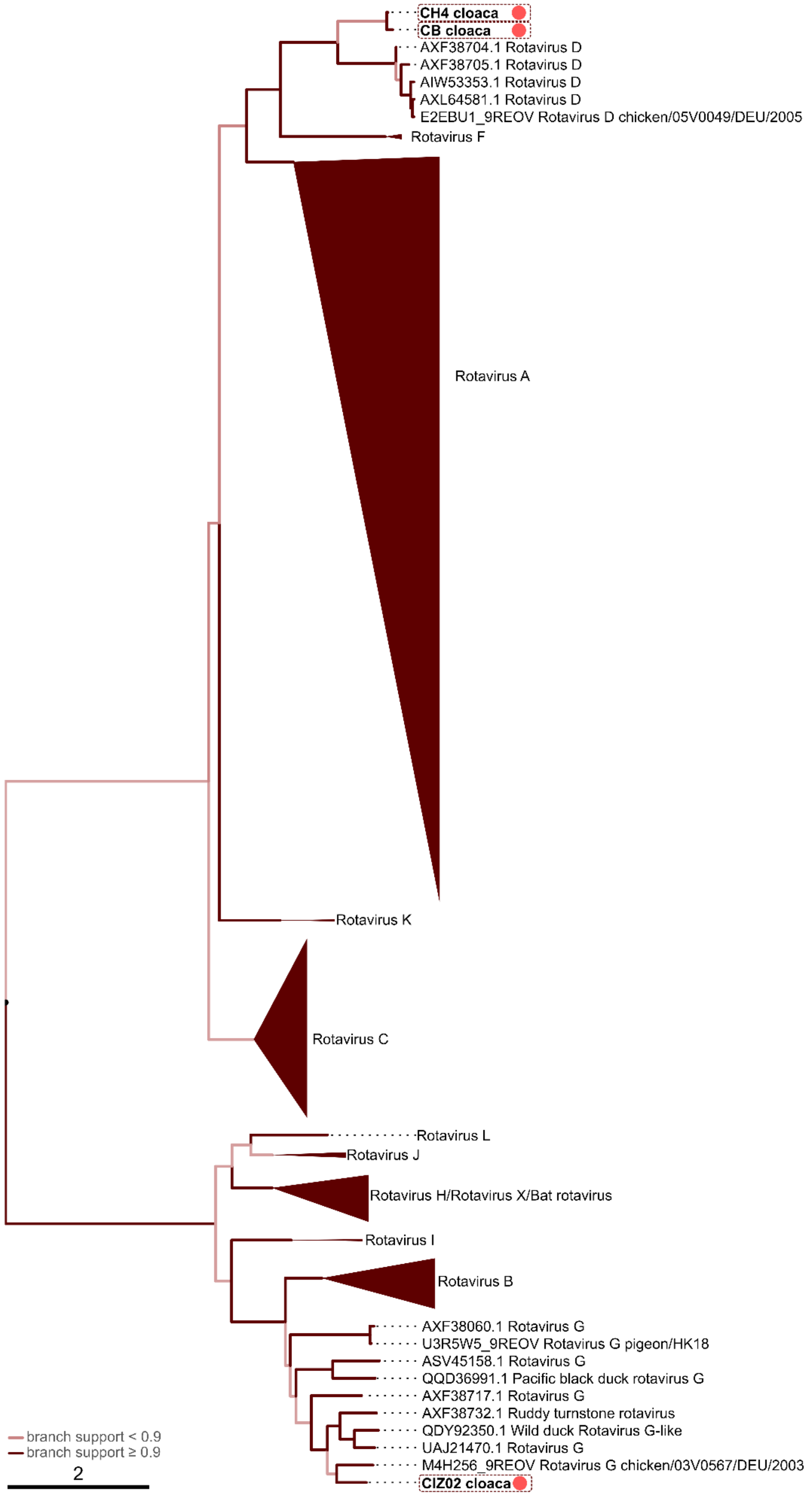


**Supplementary Figure 6. Phylogenetic tree of rotavirus VP4 (product of segment 4) sequences obtained in this study (red dots) and published rotavirus VP4 sequences**
Phylogenies based on amino acid sequences.

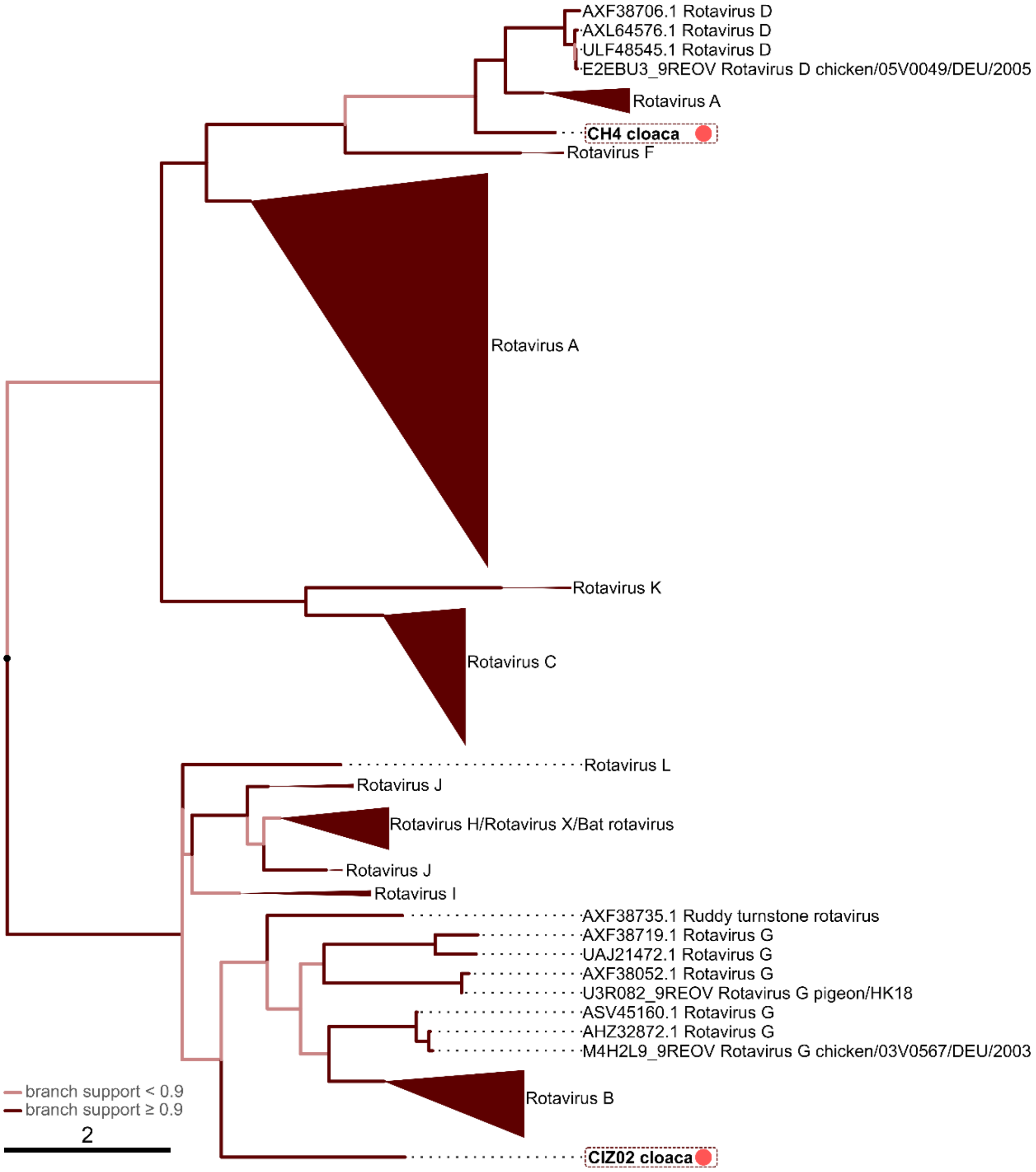


**Supplementary Figure 7. Phylogenetic tree of rotavirus NSP1 (product of segment 5) sequences obtained in this study (red dots) and published rotavirus NSP1 sequences**

Phylogenies based on amino acid sequences. Segment 5 of rotavirus B, G, and I contains two ORFs[2] (also refer to ICTV), and initial alignments were reduced to only include the likely peptide 2 subtree. That some rotavirus A sequences are more closely related to rotavirus D than other rotavirus A has been described before.[3]

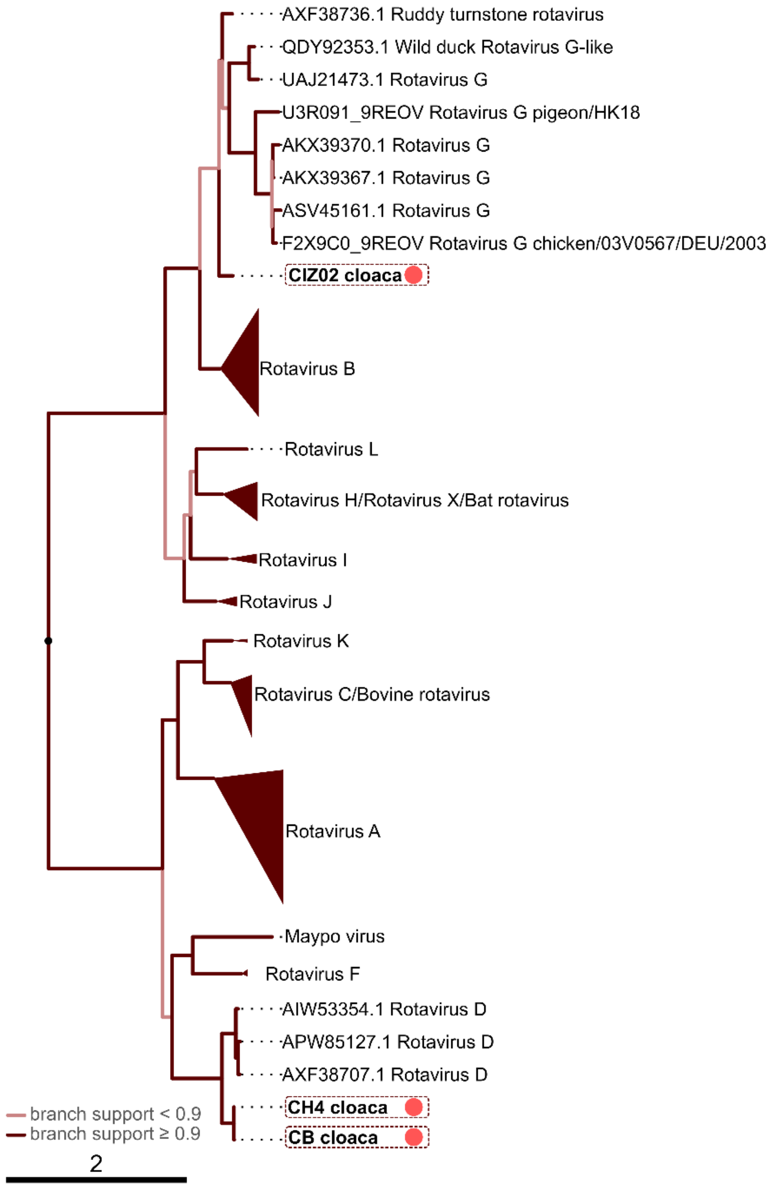


**Supplementary Figure 8. Phylogenetic tree of rotavirus VP6 (product of segment 6) sequences obtained in this study (red dots) and published rotavirus VP6 sequences**
Phylogenies based on amino acid sequences.

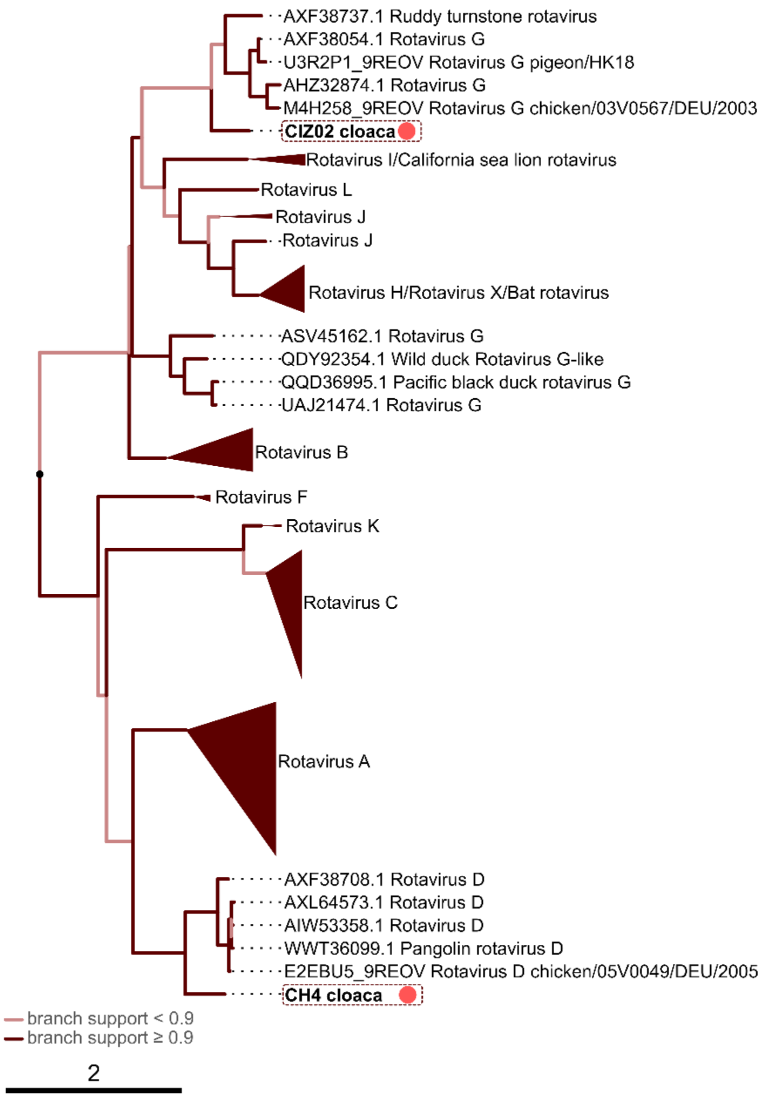


**Supplementary Figure 9. Phylogenetic tree of rotavirus NSP3 (product of segment 7) sequences obtained in this study (red dots) and published rotavirus NSP3 sequences**
Phylogenies based on amino acid sequences.

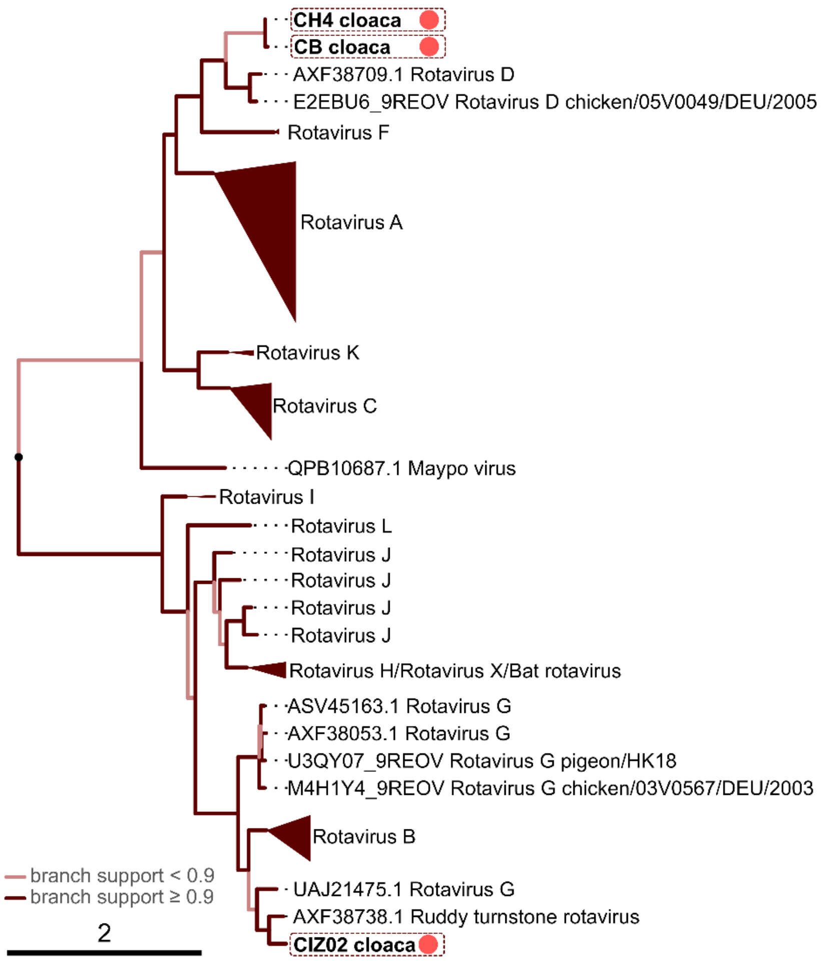


**Supplementary Figure 10. Phylogenetic tree of rotavirus NSP2 (product of segment 8) sequences obtained in this study (red dots) and published rotavirus NSP2 sequences**
Phylogenies based on amino acid sequences.

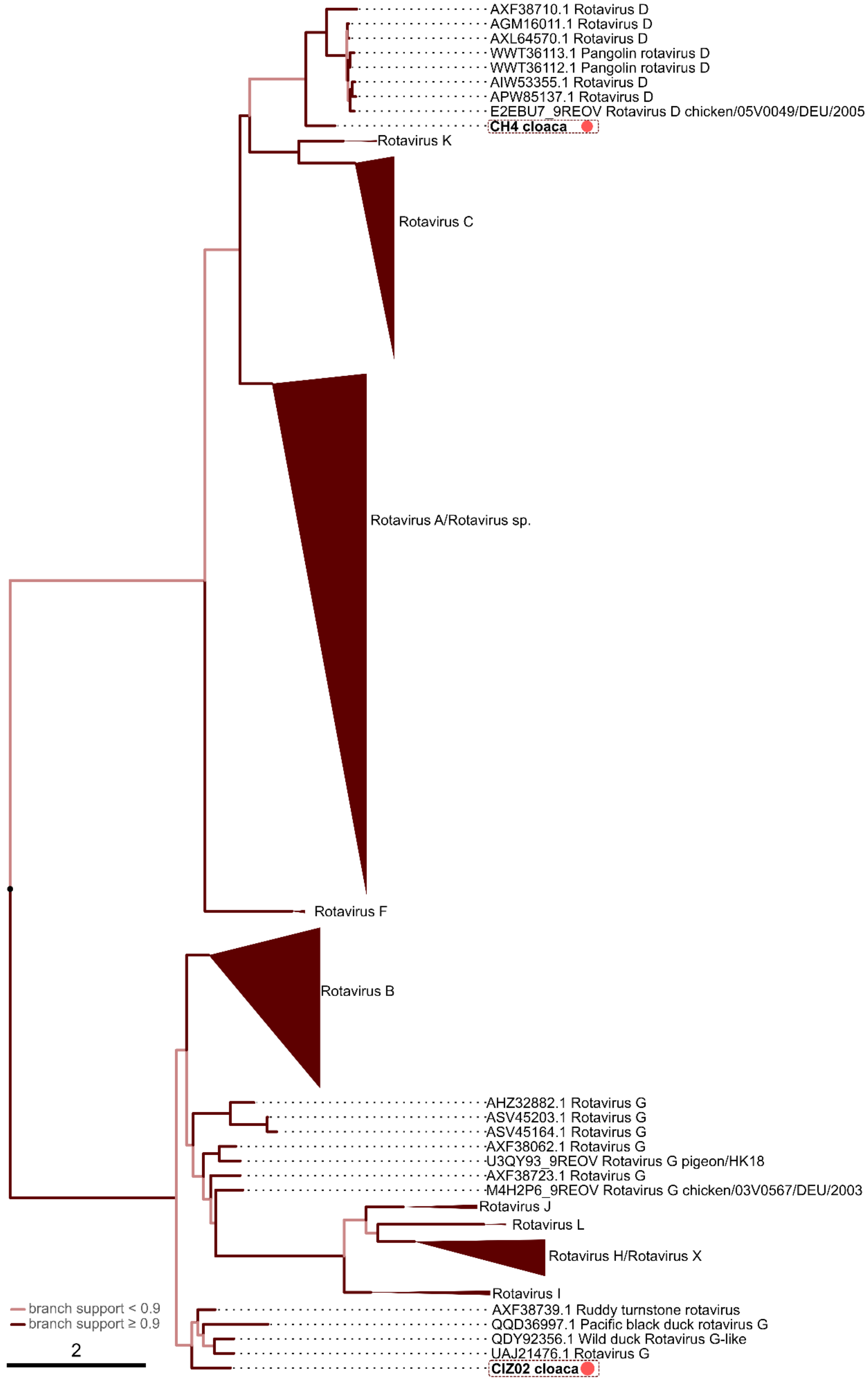


**Supplementary Figure 11. Phylogenetic tree of rotavirus VP7 (product of segment 9) sequences obtained in this study (red dots) and published rotavirus VP7 sequences**
Phylogenies based on amino acid sequences.

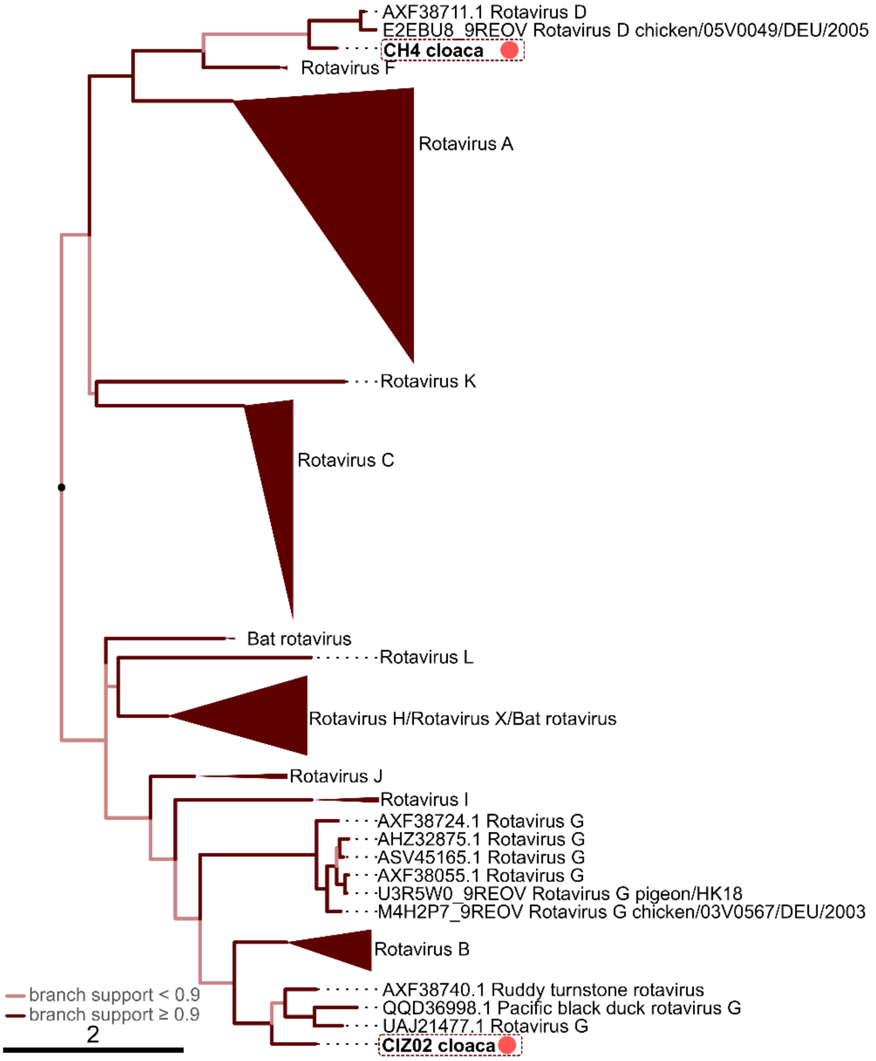


**Supplementary Figure 12. Phylogenetic tree of rotavirus NSP4 (product of segment 10) sequences obtained in this study (red dots) and published rotavirus NSP4 sequences**
Phylogenies based on amino acid sequences.

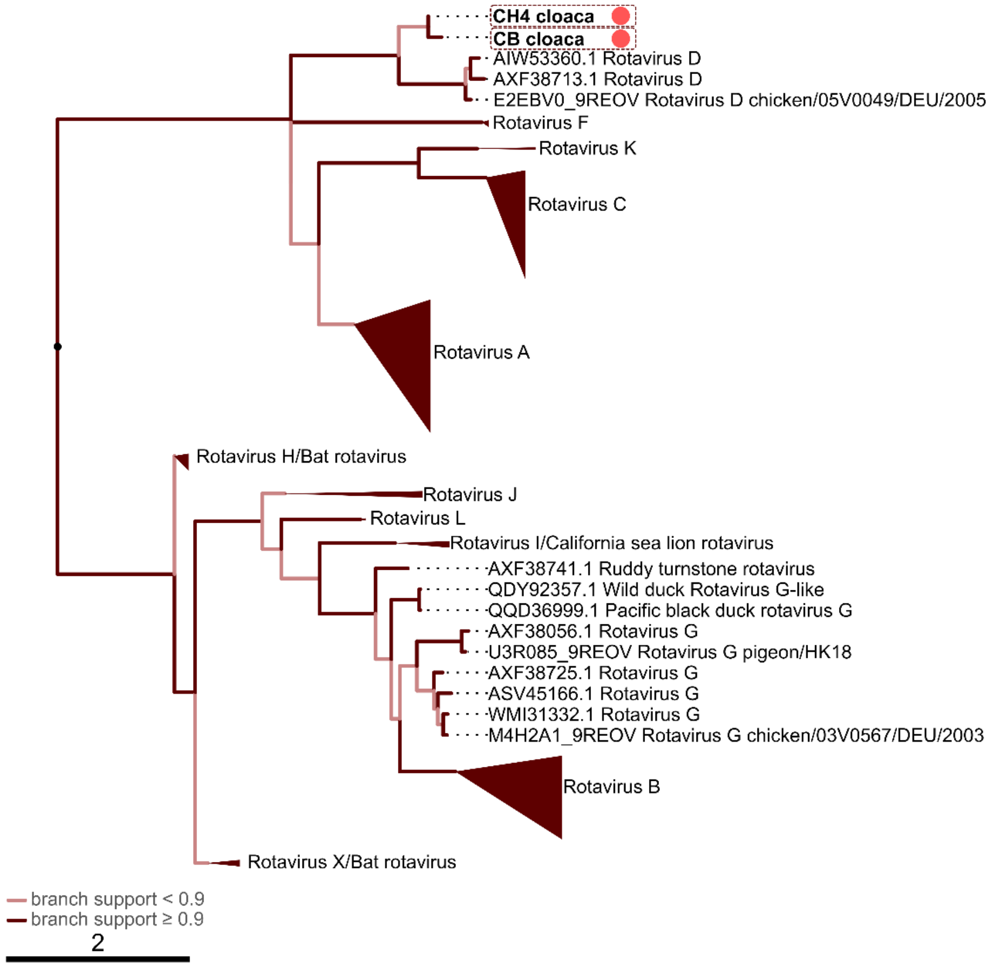


**Supplementary Figure 13. Phylogenetic tree of rotavirus NSP5 (product of segment 11) sequences obtained in this study (red dots) and published rotavirus NSP5 sequences**
Phylogenies based on amino acid sequences.

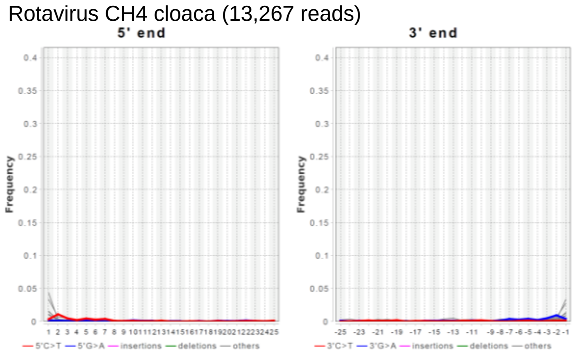


**Supplementary Figure 14. Damage patterns of reads aligned to rotavirus consensus sequences in modern *Pygoscelis adeliae* mummy RNA library reads of CH4 cloaca**

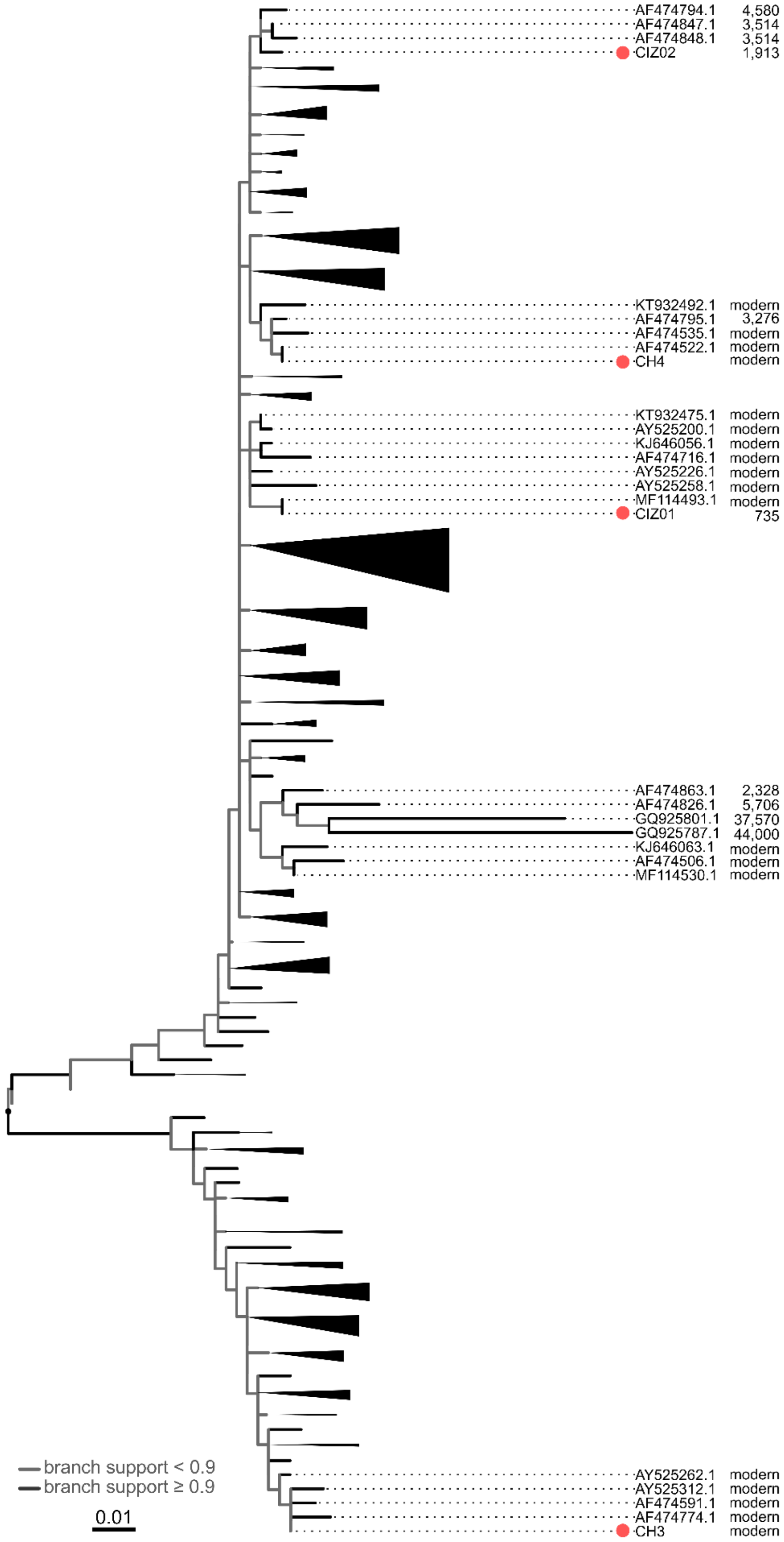


**Supplementary Figure 15. Phylogenetic tree of DNA-based Adélie penguin mitogenomes obtained in study (red dots) and published Adélie penguin mitogenomes**

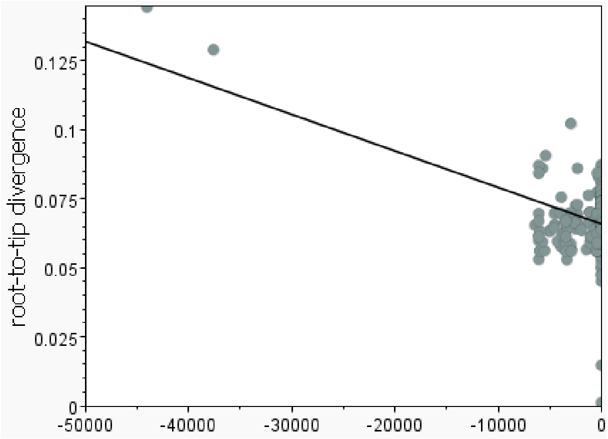


**Supplementary Figure 16: Root-to-tip divergence as function of time for Adélie penguin mitogenomes obtained in study and published Adélie penguin mitogenomes**
Sequences were outlier-filtered before calculation (see main text). $R^2$=0.20, correlation coefficient -0.44 (based on heuristic residual mean squared error function in TempEst[4]).

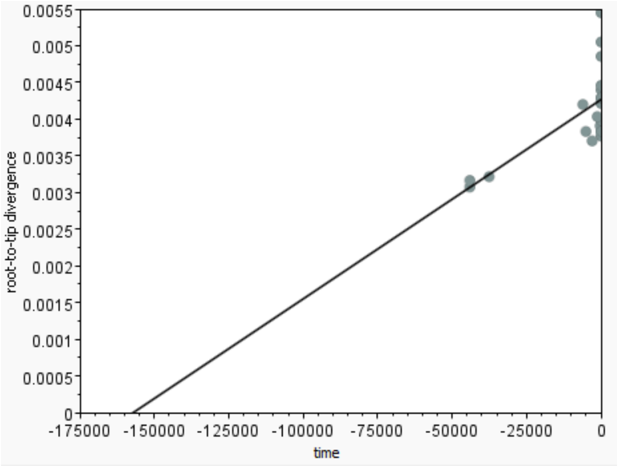


**Supplementary Figure 17. Root-to-tip divergence as function of time for Adélie penguin mitogenomes published by Subramanian et al. (2009)[5]**
$R^2$=0.47, correlation coefficient 0.68 (based on heuristic residual mean squared error function in TempEst[4]).

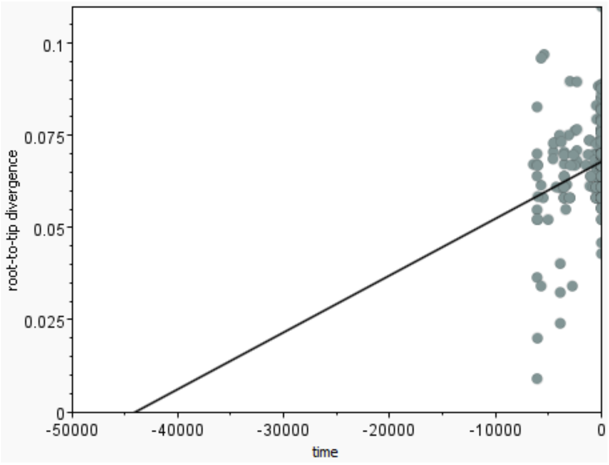


**Supplementary Figure 18. Root-to-tip divergence as function of time for Adélie penguin mitogenomes published by Lambert et al. (2002)[6]**
$R^2$=0.06, correlation coefficient 0.24 (based on heuristic residual mean squared error function in TempEst[4]).

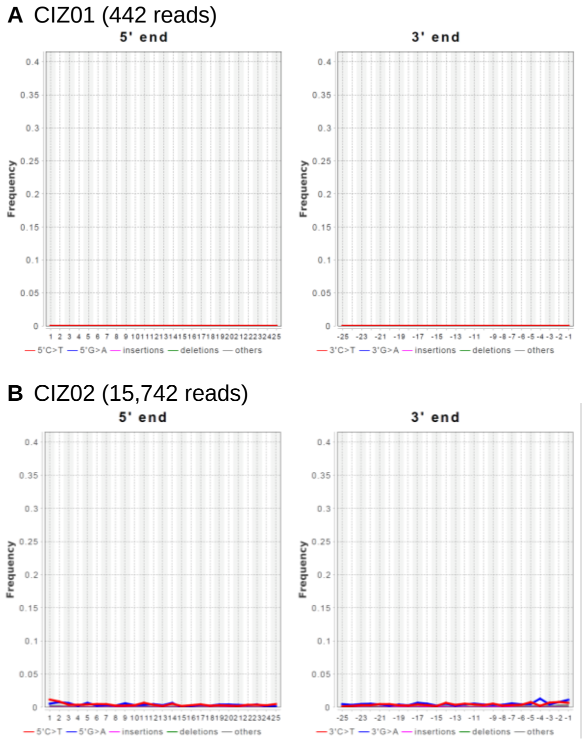


**Supplementary Figure 19. Damage patterns of ancient *Pygoscelis adeliae* mummy DNA library reads aligned to their respective consensus sequences**
(A) Adélie penguin mummy CIZ01.
(B) Adélie penguin mummy CIZ02.

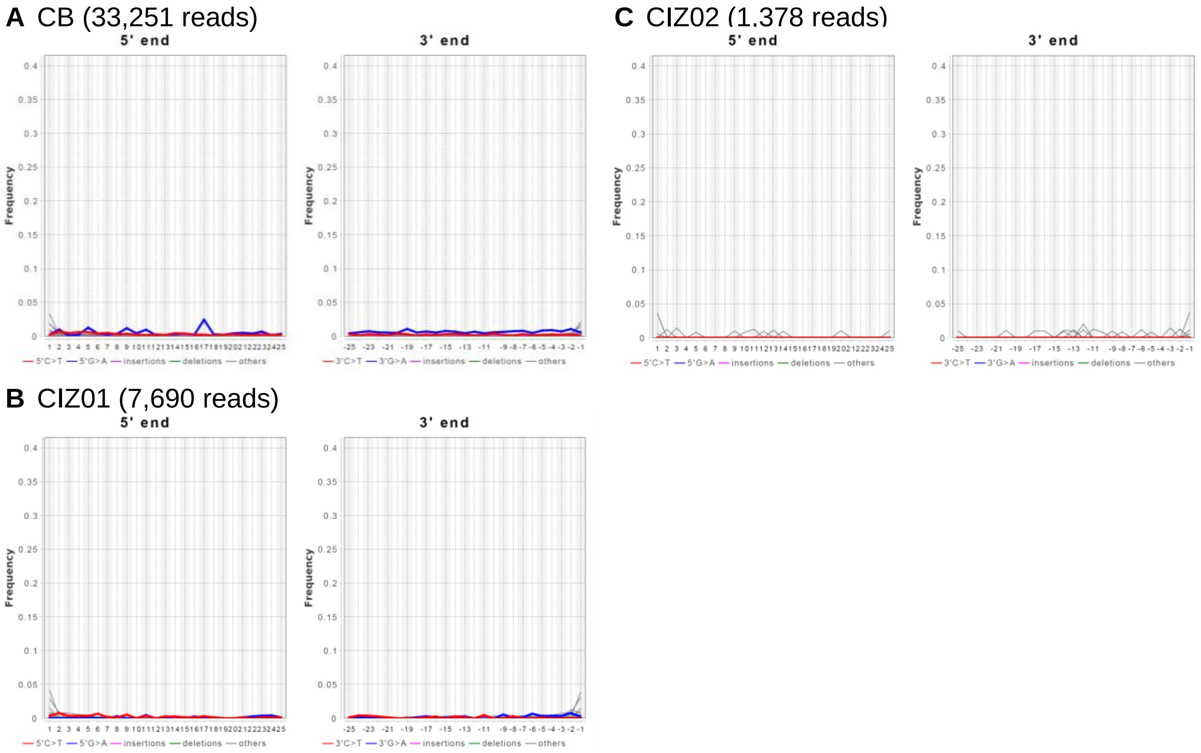


**Supplementary Figure 20. Damage patterns of ancient *Pygoscelis adeliae* mummy RNA library reads aligned to their respective RNA consensus sequences**
(A) Adélie penguin mummy CB.
(B) Adélie penguin mummy CIZ01.
(C) Adélie penguin mummy CIZ02.

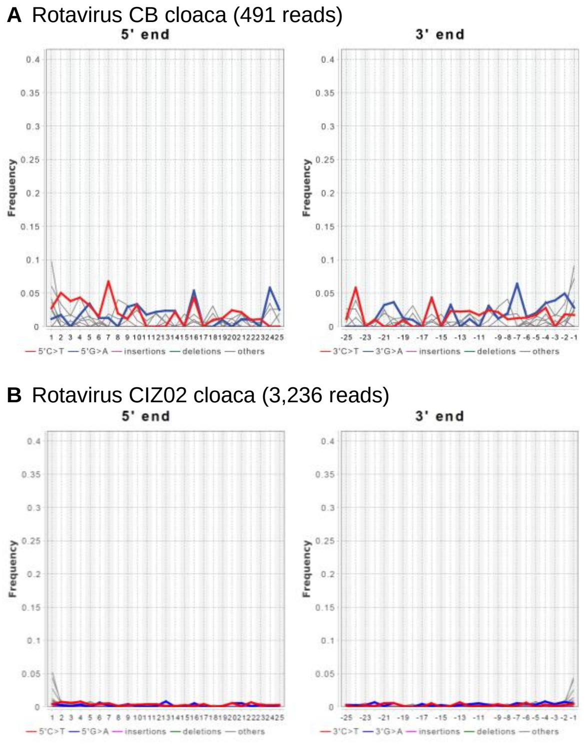


**Supplementary Figure 21. Damage patterns of reads aligned to rotavirus consensus sequences in ancient *Pygoscelis adeliae* mummy RNA library reads**
(A) Adélie penguin mummy CB.
(B) Adélie penguin mummy CIZ02.

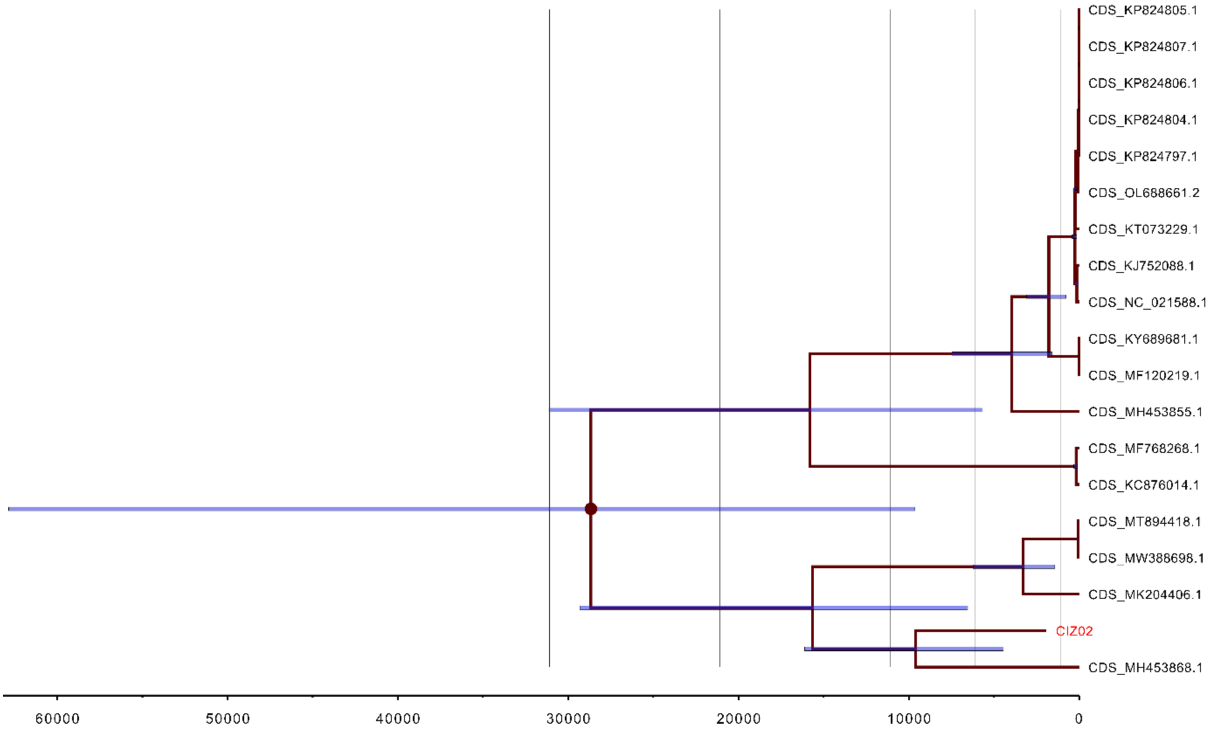


**Supplementary Figure 22. Phylogenetic tree of the CIZ02 rotavirus segment 6 sequence obtained in this study (CIZ02) and published rotavirus G segment 6 sequences**
Phylogenies based on nucleic acid sequences. Blue bars: Node error bars.

#### Supplementary Tables

**Supplementary Table 16. Results of radiocarbon dating of ancient penguin mummies**
UGAMS: University of Georgia AMS laboratory; NOSAMS: Woods Hole National Ocean Sciences AMS Facility; cal. yr. BP: calibrated years before present.

| Specimen | Laboratory number | 2 sigma cal. yr. BP date range | Mean of min/max dates (cal. yr. BP) per measurement | Mean of min/max dates (cal. yr. BP) across both replicates |
|---|---|---|---|---|
| CB | UGAMS 69031 | 431-60 | 246 | 280 |
| | NOSAMS 192400 | 499-183 | 341 | |
| CIZ01 | NOSAMS 136469 | 946-722 | 834 | 735 |
| | UGAMS 67374 | 834-524 | 679 | |
| CIZ02 | UGAMS 67375 | 2113-1724 | 1919 | 1913 |
| | UGAMS 68269 | 2106-1712 | 1909 | |

**Supplementary Table 18. Sample names, batch numbers, and sequencing platforms of the Adélie penguin mummy samples used for sequencing in this study**

| Sample | Batch Number | Sequencing Platform |
|---|---|---|
| CH1 | 2 | Element Biosciences |
| CH2 | 2 | Element Biosciences |
| CH3 | 1 | Illumina |
| CH4 | 1 | Illumina |
| CB | 2 | Element Biosciences |
| CIZ01 | 1, 2 | Illumina |
| CIZ02 | 1, 2 | Illumina |
